# A general quantitative relation linking bacterial cell growth and the cell cycle

**DOI:** 10.1101/795013

**Authors:** Hai Zheng, Yang Bai, Meiling Jiang, Taku A. Tokuyasu, Xiongliang Huang, Fajun Zhong, Xiongfei Fu, Nancy Kleckner, Terence Hwa, Chenli Liu

**Author notes:** equal contribution (Hai Zheng, Yang Bai, Meiling Jiang).

## Abstract

The foundation of bacterial cell cycle studies has long resided in two interconnected dogmas between biomass growth, DNA replication, and cell division during exponential growth: the SMK growth law that relates cell mass (a measure of cell size) to growth rate^1^, and Donachie’s hypothesis of a growth-rate-independent initiation mass^2^. These dogmas have spurred many efforts to understand their molecular bases and physiological consequences^3–12^. Most of these studies focused on fast-growing cells, with doubling times shorter than 60 min. Here, we systematically studied the cell cycle of *E. coli* for a broad range of doubling times (24 min to over 10 hr), with particular attention on steady-state growth. Surprisingly, we observed that neither dogma held across the range of growth rates examined. In their stead, a new linear relation unifying the slow- and fast-growth regimes was revealed between the cell mass and the number of cell divisions it takes to replicate and segregate a newly initiated pair of replication origins. This and other findings in this study suggest a single-cell division model, which not only reproduces the bulk relations observed but also recapitulates the adder phenomenon established recently for stochastically dividing cells^13–15^. These results allowed us to develop quantitative insight into the bacterial cell cycle, providing a firm new foundation for the study of bacterial growth physiology.

---

A fundamental notion in the quantitative study of bacterial physiology is steady state growth^16,17^, where the rate of total cell mass growth (*λ*_*m*_) is identical to the rate of cell number growth (*λ*_*c*_). To cover both the slow- and fast-growth regimes, we cultured *E. coli* K12 MG1655 cells in 32 different growth media with corresponding growth rates ranging from 0.06 to 1.7 h^−1^ (doubling times ranging from ~700 min to 24 min; see Extended Data Table 1). Special care was taken to ensure that all experimental cultures in this study lie on the steady-state line *λ*_*m*_ = *λ*_*c*_, hereafter designated as *λ*; see Extended Data Fig. 1.

The SMK growth law^1^, long-accepted in the fast-growth regime, states that the population-averaged cellular mass 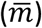 scales exponentially with growth rate, 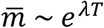, with a constant parameter *T* ≈ 1 *h*. We examined several measures of 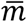, including dry weight, optical density (OD), and cell size (by flow cytometry and microscopic imaging) (**Supplementary Methods**). For each measure, we found the SMK law to break down for growth rates below 0.7 h^−1^ (Fig. 1a-d). Our OD data appear in good agreement with that obtained by Schaechter *et al.^1^* (green squares in Fig. 1e) for the mostly fast growth rates they analyzed, even though they studied a *Salmonella* strain. We also extracted cell size data from recent studies by Si *et al.^3^* and Gray *et al.^18^*, and found their data to be consistent with ours (blue and red symbols, Fig. 1f). Aside from the average cell size, even the size distributions appear to be indistinguishable for a variety of growth media^18^ (Extended Data Fig. 2a). Given the equivalence of different measures of cell mass (Extended Data Fig. 2b-d), OD_600_⋅ml per cell shall be used as the measure of 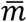 throughout the rest of this study for convenience.

**Figure 1.**
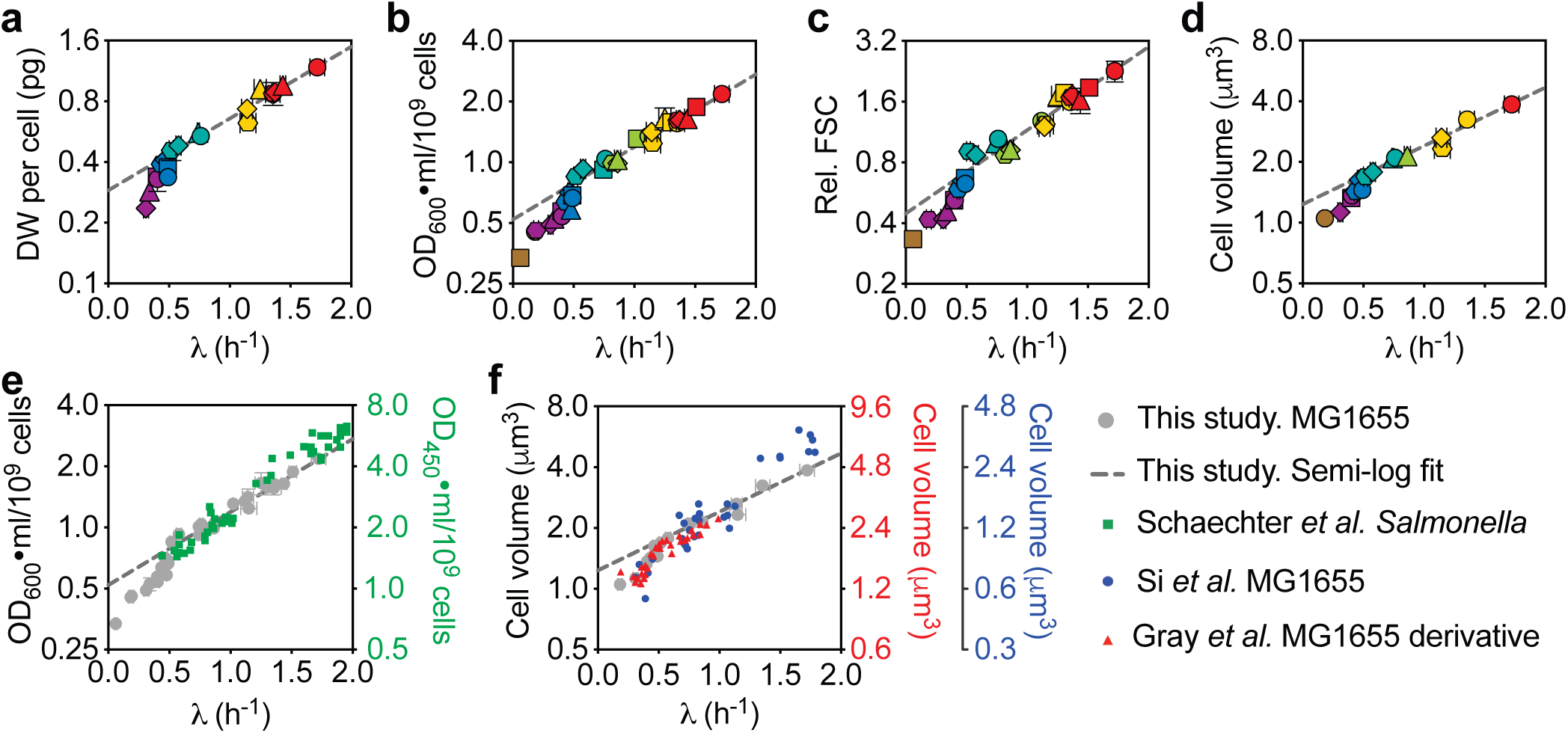
SMK growth law is violated under slow-growth conditions. **a-d**, Population-averaged dry weight (DW) per cell (**a**), OD_600_⋅ml/10^9^ cells (**b**), relative FSC (normalized by the FSC of cells grown in M18) (**c**), or cell volume (**d**) plotted against growth rate on semi-log axes. All experimental cultures were ensured to be in steady state (see **Extended Data Fig. 1** and **Supplementary Methods**). Different symbols indicate different growth media (see Extended Data Table 1 for details). Error bars in panels **a-c** represent the SDs of biological replicates (see Extended Data Table 1 for details); those in panel **d** represent 95% CI of population-averaged cell volume. Many error bars were smaller than the size of the symbols. The straight dashed lines are semi-log fit to the data in fast-growth regime (*λ* > 0.7 h^−1^); the slopes of the lines for **a-d** are 0.82, 0.83, 0.95, and 0.67, respectively. Data from slow-growth regime deviated from the lines in all cases. **e**, Comparison between the optical density data from panel **b** (gray filled circles) and the original data from Schaechter *et al^1^*. **f**, Comparison between the cell volume data from panel **d** (gray filled circles) and the data sets extracted from recent reports, Gray *et al^18^* (red triangles, 1^st^ right y-axis) or Si *et al^3^* (blue dots, 2^nd^ right y-axis). The data of Gray *et al* show a clear transition at growth rate ~0.7 h^−1^ as in the current study. The behavior of the data of Si *et al* near growth rates ~0.7 h^−1^ is difficult to discern given the sparseness of the data. We note that, even though all three studies characterized the cell volume based on microscopic images, the absolute cell volumes reported by the different studies differ about two-fold. Such systematic differences likely reflect the different image processing methods used, as reported before^32^. Gray *et al^18^* and the current study used two closely related software, Oufti^42^ and MicrobeTracker^43^, respectively, to characterize cell volume, and the resulting values are considerably more comparable with each other than with Si *et al^3^*.

We wondered if the breakdown of the SMK law in the slow-growth regime was due to the growth rate dependence of the parameter *T*. It was suggested by Donachie^2^ that *T* = *C* + *D*, where *C* + *D* is the “replication-segregation time”, i.e., the time elapsed between the initiation of a round of DNA replication to the cell division at which the corresponding sister chromosomes segregate^19^. This leads to the expectation that 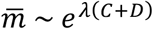 (referred to here as “Donachie’s cell-mass relation”), which was recently tested under various perturbations^3,4^. Since *e*^*λ*(*C*+*D*^) is well-established to be the population-averaged number of DNA replication origins per cell (*ō*)^20^, we measured *ō* as previously described^4,21^ (**Supplementary Methods**) across the spectrum of tested growth conditions. An exponential relation between *ō* and *λ* is seen throughout the range of growth rates (Fig. 2a), i.e.,

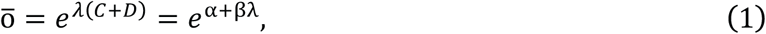

**Figure 2.**
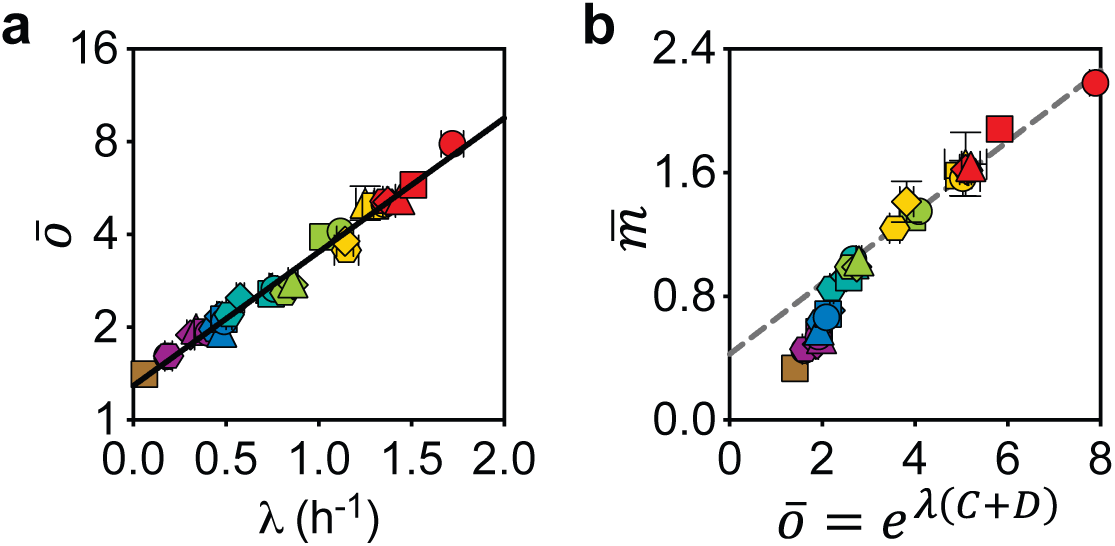
Donachie’s cell-mass relation does not hold. **a**, Correlation between the population-averaged number of DNA replication origins per cell and the growth rate. Standard run-out experiments^4,21^ followed by flow cytometry analysis were applied to quantify the *ō* (**Supplementary Methods**). The straight line is a semi-log fit *ō* = *e*^*α*+*βλ*^ (Eq. 1). **b**, Population-averaged cellular mass (in units of OD_600_⋅ml/10^9^ cells) is not proportional to *e*^*λ*(*C*+*D*)^. The straight line is linear fit to the data in fast-growth regime (*λ* > 0.7 h^−1^). The fact that the linear fit does not go through the origin clearly shows that 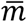 is not proportional to *e*^*λ*(*C*+*D*)^ even for fast-growth rates. Error bars represent the SDs of biological replicates (see Extended Data Table 2 for details); many of them were smaller than the size of the symbols.

with the best-fit parameters α = 0.28 and β = 0.99 h, and *R*^2^ > 0.97. Eq. (1) also leads to a Michaelis–Menten-like formula for the rate of replication-segregation process,

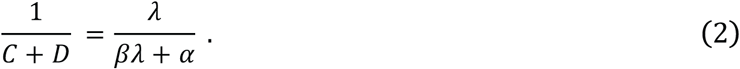

For fast growth rates where λ is much larger than *α*/*β* ≈ 0.28/*h*, this is consistent with the wide-spread notion that *C* + *D* stays roughly constant for fast growing cells^19^. A number of authors have determined population-averaged *C* + *D* or *ō* at various growth rates for *E. coli*^3,22–24^; Michelsen *et al.^23^* in particular obtained data far into the slow-growth regime. When compared to these published data, Eq. 1 provides a good fit in all cases (Extended Data Fig. 3), indicating that this simple relationship has lain undetected in such data for a considerable amount of time.

Given the validity of Eq. 1 across the growth-rate range, Donachie’s cell-mass relation would lead to 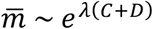, preserving the SMK growth law. However, the SMK growth law breaks down in the slow-growth regime (Fig. 1); we thus expect Donachie’s cell-mass relation to also break down. Indeed, the relation between 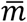 and *e*^*λ*(*C*+*D*^) (represented by *ō*) is not linear (Fig. 2b). A linear relation certainly can be fitted to the data in the fast-growth regime (dashed line in Fig. 2b). The nonzero intercept, however, indicates that Donachie’s cell-mass relation does not hold even in the fast-growth regime where the SMK law holds.

This led us to directly examine Donachie’s hypothesis on the growth-rate-independent initiation mass^2^, 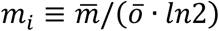^25^, which is supported by multiple recent reports^3,5,26^ and is canonically accepted in numerous modeling studies^3,27^, although challenged in some early studies^7,8,28–30^. Our data (Fig. 3a) clearly shows that the initiation mass is not constant, rather exhibiting a non-monotonic relation over growth rates with a peak around *λ* ≈ 0.7/*h* (doubling time of 1 hr). Our data are mostly consistent with those from recent studies, although the sparsity and variability of those data gave the appearance of a constant initiation mass: Wallden *et al.^26^* directly measured the single cell initiation volume in three different growth conditions and found it to be constant. However, we note that the conditions tested in their study were at different temperature (Fig. 3b), which is known to affect growth rate but has little effect on cell mass^1^, making it difficult to compare to our results obtained at 37 °C. Si *et al.^3^* used the combination of microscopic imaging and run-out experiments to quantify the initiation volume. They reported the constancy of initiation mass for different growth rates under various perturbations. Their data however shows considerable variability (green diamonds, Fig. 3c) and cannot rule out the tight trend suggested by our data (grey circles).

**Figure 3.**
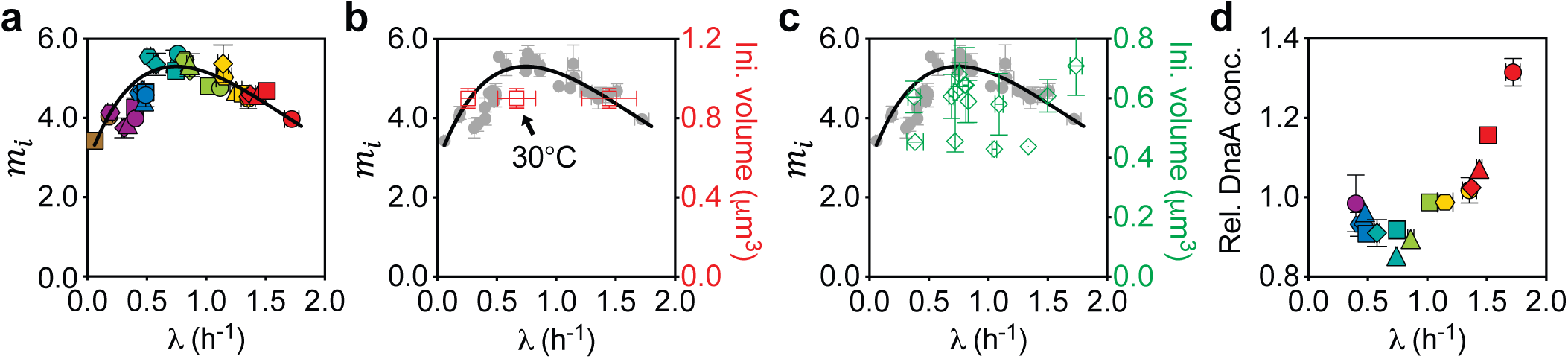
Donachie’s constant-initiation-mass hypothesis breaks down. **a**, The initiation mass exhibits a non-monotonic dependence on growth rate. The initiation mass *m*_*i*_ (in units of 10^−10^ OD_600_⋅ml) for each growth condition was characterized as described in the main text. It varied by ~50 % across different growth rates, reaching a peak value ~5.5 × 10^−10^ OD_600_⋅ml at a growth rate 0.7 h^−1^. The growth rate dependence of the initiation mass is well described by the black curve (Eq. 4). **b**, Comparison between the data from panel **a** (gray filled circles) and those extracted from Wallden *et al*^26^ (MG1655, red squares, right y-axis). For Wallden *et al*’s data, the error bars for initiation volume are set to 10 % as they described; the error bars for growth rate represent the SDs at the single cell level as they described. The culture temperature for the middle data point (arrow) is 30 °C, while the other two data points and ours are obtained at 37 °C. **c**, Comparison between the data from panel **a** (gray filled circles) and those extracted from Si *et al^3^* (MG1655, green diamonds, right y-axis). The error bars of Si *et al*’s data represent the SDs of biological replicates. **d**, Relative DnaA protein concentration exhibits a non-monotonic dependence on growth rate. Relative DnaA protein concentration varied by ~ 50%, reaching a minimum value at ~ 0.7 h^−1^, approximately where the initiation mass reaches a peak. It was quantified by quantitative proteomics and normalized by the population average of the total protein concentration across growth rates (see **Supplementary Methods**). Error bars in panels **a** or **d** represent the SDs of biological replicates (see Extended Data Table 2 for details); many of them were smaller than the size of the symbols.

At the molecular level, we note that DnaA is centrally involved in the initiation of DNA replication in *E. coli*. Its expression level was found to correlate negatively with the initiation mass in a titratable strain^6^. Using quantitative proteomics (**Supplementary Methods**), we found that the DnaA protein concentration negatively correlated with the initiation mass in wild type cells (Extended Data Fig. 4), and their growth-rate dependences were mirror images of each other (compare Fig. 3d, 3a). This anti-correlation does not necessarily imply causation but could be the result of an underlying mechanism that controls both.

The breakdown of the two longstanding dogmas might suggest different mechanisms governing cell mass in the slow- and fast-growth regimes. Nevertheless, directly plotting the data in Fig. 2b against λ(*C* + *D*) revealed a crisp, linear relation (*R*^2^ > 0.97) across the entire growth-rate range examined (Fig. 4a),

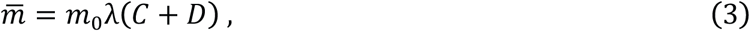

**Figure 4.**
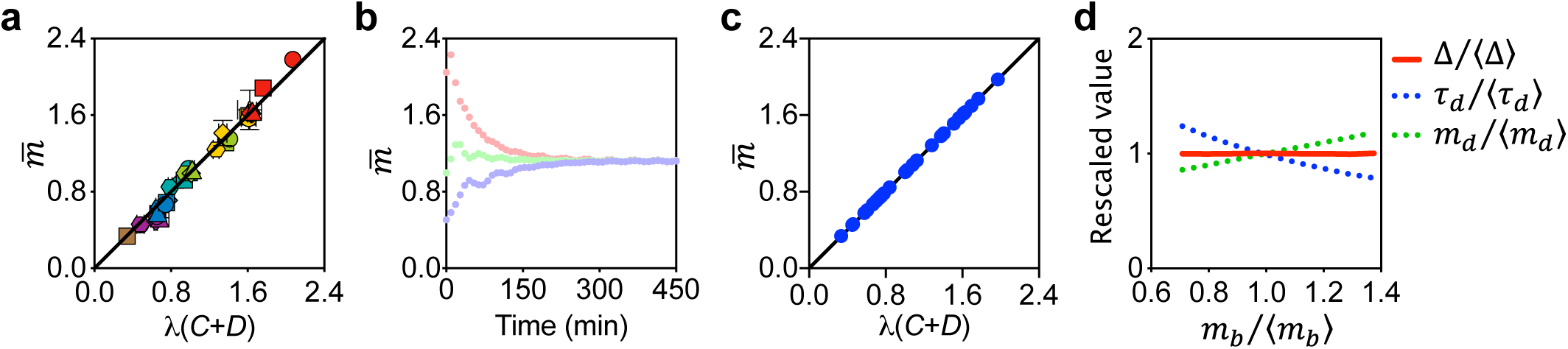
New linear relation unifies the slow- and fast-growth regimes. **a**, The population-averaged cellular mass scales linearly with *λ*(*C* + *D*), a measure of the generation number during the time from initiation of a round of DNA replication to the cell division at which the corresponding sister chromosomes segregate^31^. Error bars represent the SDs of biological replicates (see Extended Data Table 2 for details); many of them were smaller than the size of the symbols. **b**, Simulations of the ‘integral-threshold model’ (see **Supplementary Models**) showed that the population-averaged cellular mass approaches the expected steady-state value independent of the initial conditions. Three different initial conditions were used [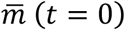 equals to 2.0, 1.0, or 0.5]. **c**, Simulations reproduced the linear relation (Eq. 3), where the population-averaged cellular mass is calculated by ln(2)⋅〈*m*_*d*_〉. **d**, Simulations captured the adder phenomenon. The normalized added-mass between cell birth and division (Δ/〈Δ〉) is independent of the normalized mass at birth (*m*_*b*_/〈*m*_*b*_〉), whereas the normalized mass at division (*m*_*d*_/〈*m*_*d*_〉) or the normalized interdivision time (*τ*_*d*_/〈*τ*_*d*_〉) are positively or negatively correlated with the normalized mass at birth, respectively. The related correlation coefficients were 0.51 and 0.49, which are in agreement with the experiment measurements^15^. The growth rates in panel **b** and **d** were set as 0.92 h^−1^. Similar results were observed across entire spectrum of growth rates (data not shown).

with the best-fit slope *m*_0_ = 1.00 ± 0.013 OD_600_⋅ml/10^9^ cells or 0.545 ± 0.015 pg DW/cell (Extended Data Fig. 2b). This simple relation suggests the existence of a unifying mechanism governing cell cycle progression in both slow- and fast-growth conditions. The population-averaged cellular mass is proportional to λ(*C* + *D*), which is a measure of the number of generations elapsed during the period of *C* + *D*^31^.

Eqs. 1 and **3** yield a growth-rate-dependent initiation mass,

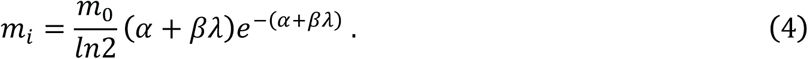

This relation peaks at α + βλ = *λ*(*C* + *D*)= 1 (doubling time ~55 min) and captures well the non-monotonic behavior of our experimental data (solid line in Fig. 3a).

The newly established cell-mass relation, Eq. 3, can be used to infer possible rules underlying cell division in different growth conditions. Solving for the growth rate *λ* in Eq. 3 and using the exponential increase in cell number, *N*(*t*) ∝ *e*^*λt*^, we have 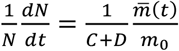. Since the left-hand side of this equation is just the rate of cell-number doubling, the form of this equation prompts a *single-cell* interpretation of the stochastic cell division dynamics: The right-hand side can be interpreted as the (probabilistic) rate of cell division, with the single cell mass *m*(*t*) ~ *e*^*λt*^. This simple stochastic cell division model reproduces **Eq. 3** and other properties (Extended Data Fig. 5a-c), however does not capture the distribution of division mass and inter-division time (Extended Data Fig. 6a-c), due in part to the non-zero probability in the model for a second division to occur immediately following a first one (**Supplementary Models 2-3**). This prompted us to formulate a modified model (detailed in **Supplementary Models 4**), in which the probability of cell division at a time *t* since the most recent division is a function of

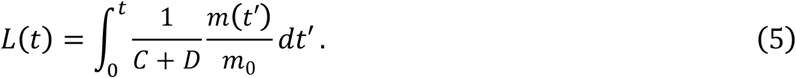

In a simple stochastic realization in which *L*(*t*) is subjected to an accumulation noise and cell division occurs when *L*(*t*) reaches a growth-rate independent threshold^32,33^, the observed distribution of division mass is obtained (Extended Data Fig. 6d-f). Simulations using this model, with the λ-dependent form of 1/(*C* + *D*) (Eq. 2), show that cells reach the same average steady-state mass regardless of the initial mass (Fig. 4b). Further, the population-averaged steady-state cellular mass is *m*_0_*λ*(*C* + *D*) across growth rates, so that Eq. 3 is reproduced (Fig. 4c). This integral-threshold cell division model also reproduces the “adder” behavior^13–15^, wherein the added mass during one cell cycle is independent of birth mass, while the inter-division time decreases and the division mass increases with increasing birth mass, respectively (Fig. 4d).

The distinct structure of this integral-threshold model suggests an underlying mechanistic basis (Extended Data Fig. 7). In brief, the quantity *L*(*t*) can be interpreted as a product of an accumulation process that licenses cell division upon reaching a threshold; this licensing product could be the divisome^34,35^, the septum^35,36^, and/or the proposed Progress Control Complex^9^, all of which are constructed prior to cell division (Extended Data Fig. 7a). 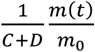 can then be interpreted as the synthesis flux, provided by one or a collection of proteins, whose abundances are proportional to cell mass independent of the growth rate (see Extended Data Fig. 7b for some candidates suggested from proteomics), and whose specific rates are proportional to 1/(*C* + *D*). We further determined the *C*-period using a deep sequencing version of standard methods^4,20^ (Extended Data Fig. 8, **Supplementary Methods**), and found 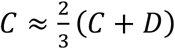 (Extended Data Fig. 7c), which implies that 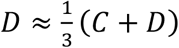. The synthesis rate of *L* could thus reflect the dynamics of chromosome replication, segregation, cell division, or their combinations. Despite the success of the integral-threshold model in describing cell mass in steady state, we note that it does not fully capture the ‘rate maintenance’ phenomenon^37^ when applied naively to growth transitions (Extended Data Fig. 9, **Supplementary Models 6**). As ‘rate maintenance’ has been used to support the link between chromosome replication and cell division^38^, a more complete model is necessary to take into account the direct influence of chromosome events on cell division^10,11^.

In this study, our experimental results from steady-state batch culture clearly demonstrated that the two long-standing dogmas, SMK growth law and Donachie’s constant-initiation-mass hypothesis, do not hold. Instead, we revealed a new linear relation between cell mass and the replication-segregation cycle, *λ*(*C* + *D*), covering both the slow- and fast-growth regimes in a unified manner. Disproving Donachie’s hypothesis has strong repercussions for existing cell-cycle models, many of which take the constant initiation mass as a central assumption^3,27,39,40^. Of the few models not relying on this hypothesis^32,33,41^, our model is the closest to that of Si *et al*^33^. which however does not capture the growth-rate dependence of cell mass (**Supplementary Models 5**). The integral-threshold model is the only model which reproduces key features of cell division at both the population and single cell levels, while also exhibiting the ‘adder phenomenon’ across growth rates. Altogether, the newly discovered quantitative relation and its associated model lay down a new basis for understanding bacterial cell cycle control.

## MATERIALS AND METHODS

Details of bacterial strains and culture conditions and methods are given in **Supplementary Methods**, and the mathematical models are in **Supplementary Models**.

## ACKNOWLEDGEMENTS

The authors acknowledge numerous colleagues for insightful discussions. This work was financially supported by National Key R&D Program of China (2018YFA0902701), Major Research Plan of the National Natural Science Foundation of China (91731302), Strategic Priority Research Program (XDPB0305) and the Key Research Program (KFZD-SW-216) of the Chinese Academy of Sciences, Shenzhen Grants (JCYJ20170818164139781, KQTD2015033117210153, Engineering Laboratory [2016]1194) to C.L., the National Natural Science Foundation of China (31700045) to H.Z., the National Natural Science Foundation of China (11804355), the Guangdong nature science foundation (2018A030310010) to Y.B, the National Natural Science Foundation of China (31570095) to X.F., Grant # NIH R01 GM025326 to N.K., and Grant # NIH R01 GM095903 to T.H.

## AUTHOR CONTRIBUTIONS

C.L. initiated and directed the research. H.Z. and M.J. carried out most of the experiments with contributions from C.L., X.H., and F.Z.. Y.B., X.F. and TH conceptualized the integral-threshold model and carried out the numerical simulations and mathematical analysis with contributions from T.T.. All authors analyzed the results and wrote the manuscript.

## COMPETING INTERESTS

The authors declare no competing interests.

## DATA AVAILABILITY

The data that support the findings of this study are available from the corresponding author upon reasonable request.

## CODE AVAILABILITY

Simulation data can be generated with the custom-made code and the parameter sets provided. The code that was used for the simulations in this study are available from the corresponding author upon reasonable request.

---

**Extended Data Figure 1.**
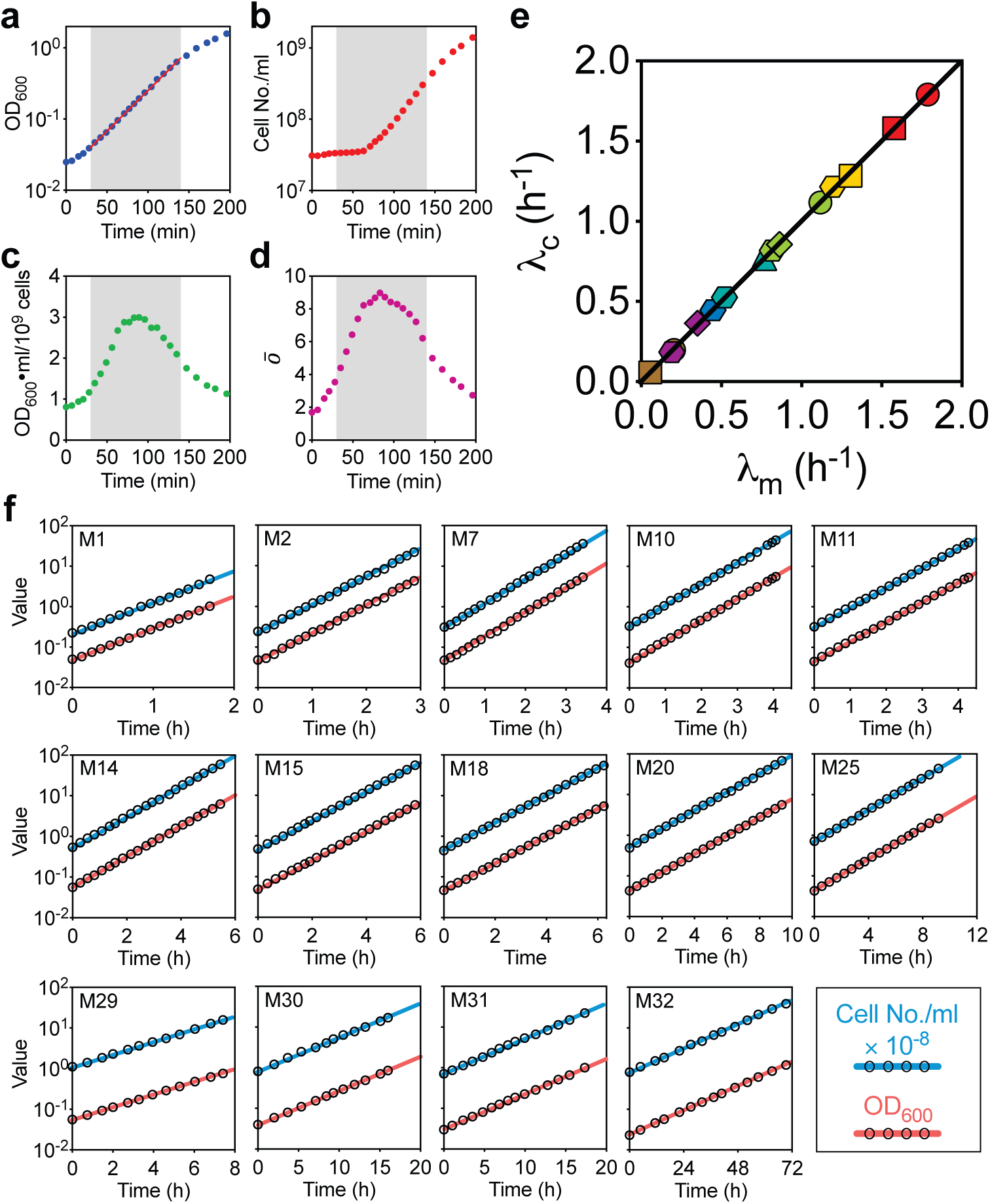
Steady-state growth validated by monitoring mass and cell-number growth simultaneously. A fundamental but often ignored point in bacterial physiology studies is the establishment of steady state growth^16,17^, where the total cell mass growth rate λ_m_ is identical to cell number growth rate λ_c_. In an exploratory experiment on early growth of cells in medium M1 (Extended Data Table 1) after inoculation from seed culture, overnight cultured cells were inoculated into pre-warmed medium with starting OD_600_ at 0.02, then grown without further dilution. The OD_600_, cell number, and population-averaged cellular origin number were characterized at different time points (**Supplementary Methods**). We found that the total cell mass growth (**a**) quickly entered exponential phase (the grey area, from 30 to 140 minutes), but cell number growth (**b**) showed a considerable lag. As a result, the average cellular mass (**c**) and origin number (**d**) varied throughout the exponential growth phase, which clearly indicated that the cells were not in true steady state. By employing serial dilutions (see **Supplementary Methods**), we found that cells grown for more than 10 mass doublings after inoculation from seed-culture were safely in steady state. This was a key step in ensuring the validity of the findings presented in this study. Following this protocol, we show in **(e)** that experimental cultures in 14 representative growth media covering the entire range of growth rates examined in this study lie on the steady state line *λ*_*m*_ = *λ*_*c*_. Representative growth curves in the steady state are shown in (f): After 10 mass doublings after inoculation from seed-culture, OD_600_ and the cell number concentration are plotted versus time, taking the dilution ratio into account to plot the “sawtooth” behavior as a single smooth curve. The cell number (red lines) and cell mass (blue lines) growth curves formed two parallel lines in semi-log plots, indicating the steady-state growth had been achieved.

**Extended Data Figure 2.**
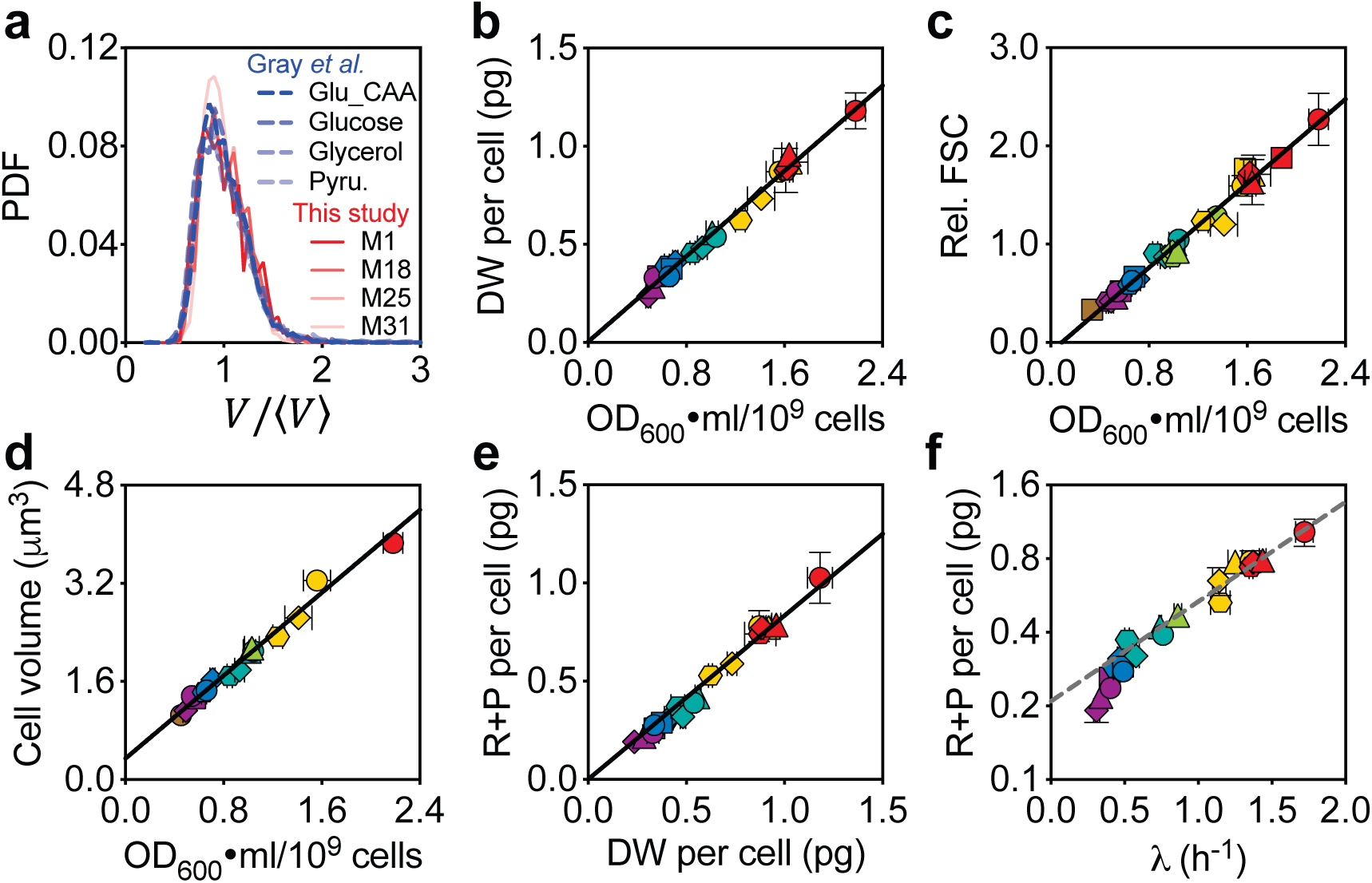
Tight correlation of different measures for cellular mass or size. **a**, Density distributions for cell volume normalized by average cell size, as quantified by automated image analysis (**Supplementary Methods**), for cells taken from the conditions described in Fig. 1d. Distributions for cells at comparable growth rates from Gray *et al^18^* were taken for comparison. When normalized by mean cell size, the different distributions appear very similar. **b-d**, The dry weight (DW) per cell (**b**), relative FSC (forward scatter) (**c**), and cell volume (**d**) plotted against the OD_600_⋅ml per 10^9^ cells. All three measures are linearly correlated with the OD_600_⋅ml per 10^9^ cells. The cell volume is expected to be a precise quantitative measure of cell size. However, data sets from different published studies^3,14,18,32^ show ~ two-fold difference for the same strain or closely related stains under similar growth conditions, possibly due to the difficulty in quantifying the actual cell diameter based on microscopic images. Given the variability in the measured FSC or cell volume, and the convenience and robustness in quantifying the cell number concentration and OD_600_, we used the OD_600_⋅ml per 10^9^ cells as the population-averaged cellular mass (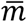) for the rest of the current study. We also measured the combined mass of RNA and protein per cell (**e**). Together, RNA and protein account for roughly 80% of total dry mass in all conditions. Plotted against growth rate **(f)**, the combined mass of RNA and protein closely resemble the trends exhibited by other measures in Fig. 1a-d.

**Extended Data Figure 3.**
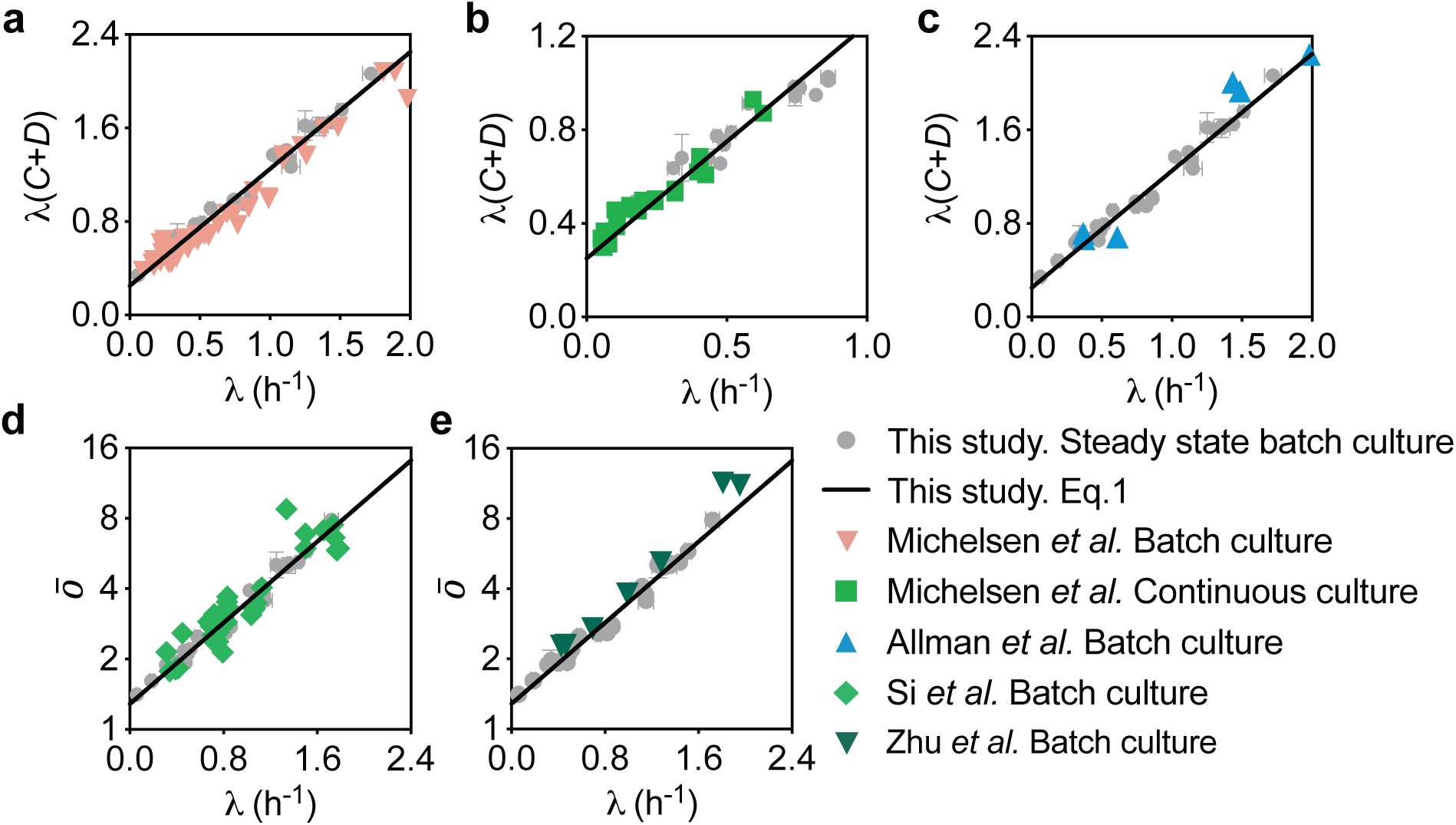
Linear relation between *λ*(*C* + *D*) and growth rate. Comparison between the data in this study and those extracted from Table 3 (**a**) or 4 (**b**) in Michelsen *et al^23^*. **c**, Comparison between the data in this study and those extracted from Allman *et al^22^*. The *C* and *D* periods were characterized by resolving the DNA histogram of cells in batch culture (**a**, **c**) or continuous culture (**b**). **d-e**, The semi-log relationship between *ō* and growth rate. **d**, Comparison between the data in this study and those extracted from Si *et al^3^*. Their *ō* were characterized by using run-out protocol followed by Hoechst 33342 staining and microscopic image analysis^3^. **e**, Comparison between the data in this study and those extracted from Zhu *et al^24^*. Their *ō* data were derived from replication origin per genome, and genome equivalents per cell for cells in batch culture. The straight lines represent **Eq.1** (**d,e**) or its derived form (**a-c**). Wallden *et al*.^26^ previously obtained three-parameter quantitative fits for the variation of *C* + *D* with the growth rate of single cells, in one to three different growth media. Eq. 1 by contrast is a single two-parameter formula that describes the relationship between *C* + *D* and λ across a wide variety of growth media, in multiple independent population-level datasets.

**Extended Data Figure 4.**
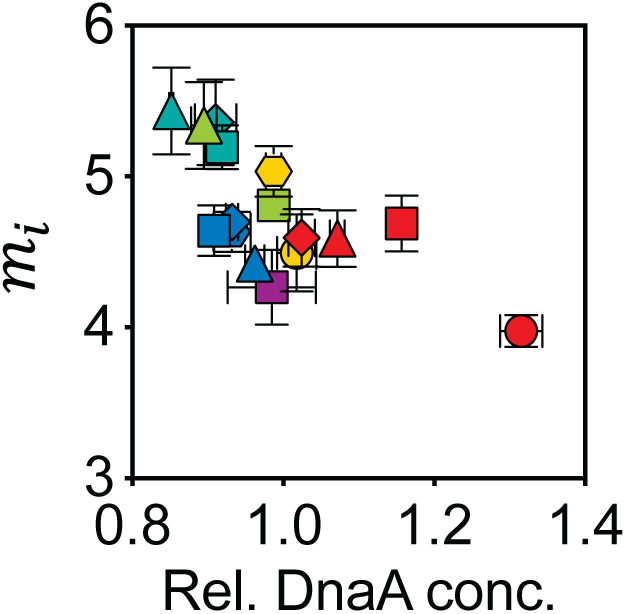
Negative correlation of initiation mass with relative DnaA protein concentration. Shown are the initiation mass from Fig. 3a and the corresponding relative DnaA concentrations from Fig. 3d. Error bars represent SDs of biological replicates (Extended Data Table 2).

**Extended Data Figure 5.**
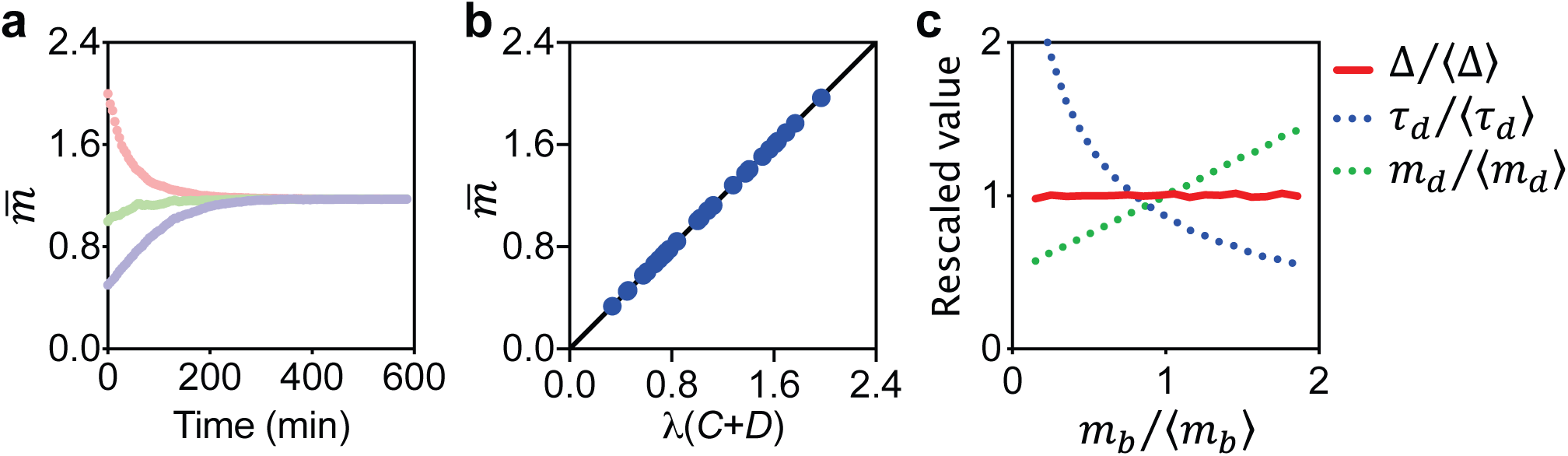
Simulation results of the ‘stochastic division model’ (see **Supplementary Models.**) **a**, The population-averaged cellular mass approached the expected steady-state value independent of the initial conditions 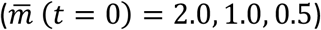. **b**, Simulations reproduced the linear relation (Eq. 3). **c**, Simulations captured the adder phenomenon. The growth rates in panel **a** and **c** were set as 0.92 h^−1^. Similar results were observed across entire spectrum of growth rates (data not shown).

**Extended Data Figure 6.**
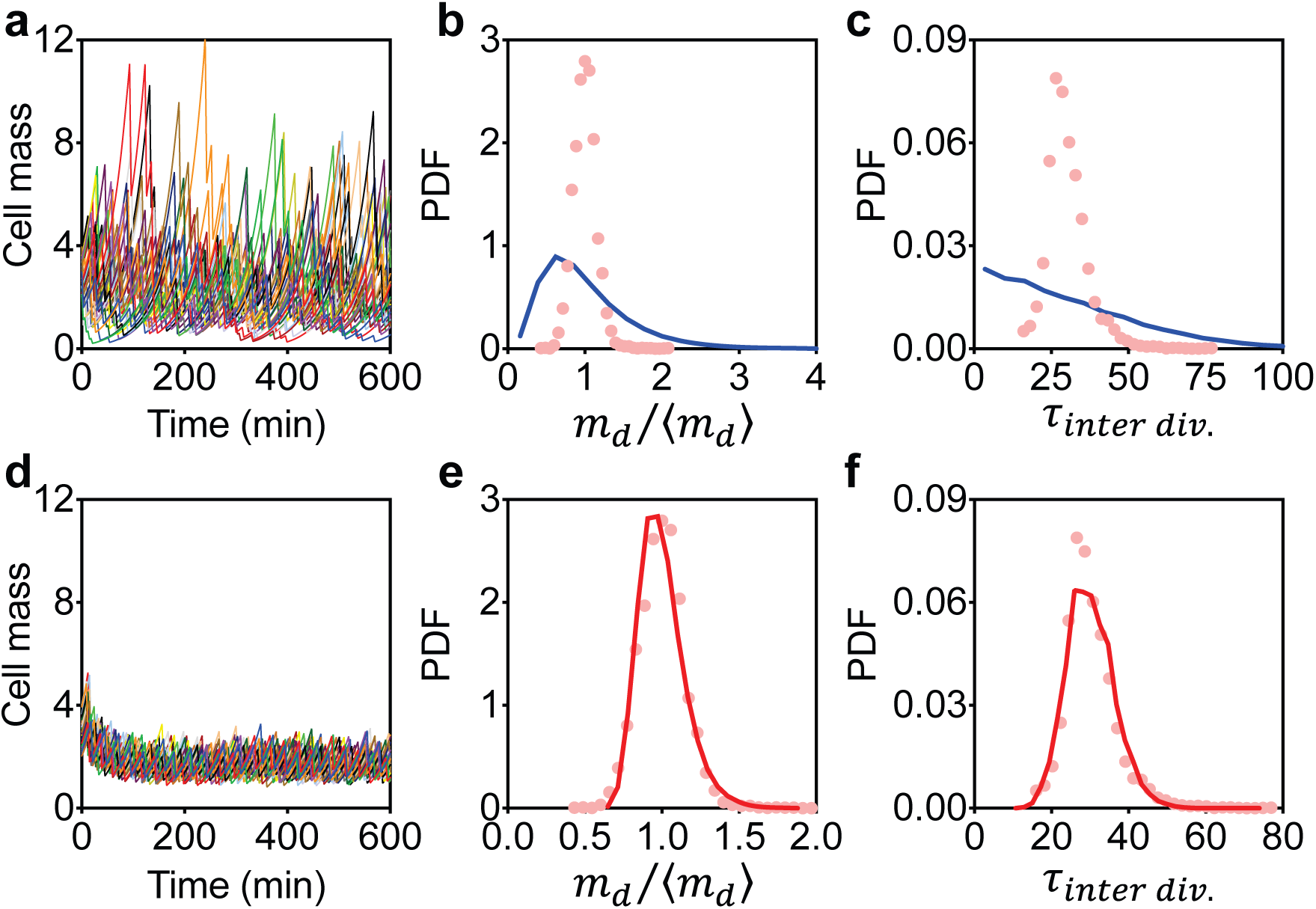
The division mass and inter division time distributions. **a,d** Simulated single cell mass dynamics. Each line represents one simulated cell with different initial cell mass (30 cells total). **b,c,e,f** Normalized cell division mass and inter-division time distributions were compared between simulations (lines) and the experimental data from Wallden *et al*^26^ (data of cell volume is regarded as cell mass). Simulations were performed in the mother machine logic where only one cell is kept after division. Simulation results were obtained when the average cell mass reached steady state. Panels **a-c** represent model predictions of the stochastic cell division model, while **d-f** represent model predictions of the integral threshold model.

**Extended Data Figure 7.**
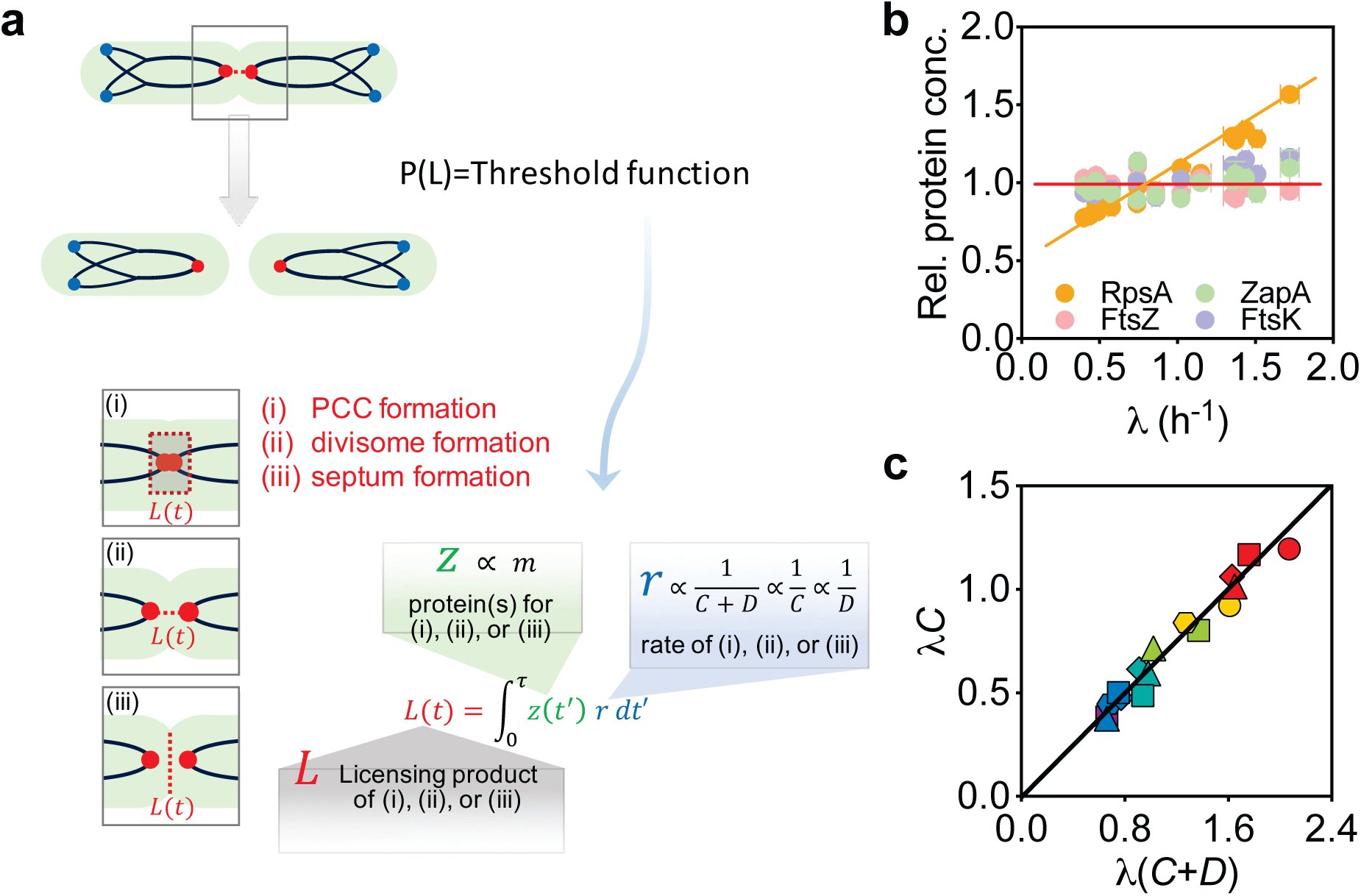
Hypothetic mechanistic basis of the integral threshold model. **a**, The key feature of our cell-division model is that the probability of cell division depends on the synthesis of a licensing product, which could be (i) progression control complex^9^, (ii) divisome^34,35^, (iii) septum^34,35^, or their combinations. The synthesis of this product, accumulated to a level *L*(*t*) at time *t* since the last division and reset to zero at division, can be implemented by one or a collection of proteins of abundance *z* per cell and with the specific rate *r*, such that 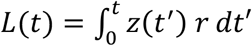. To reproduce the newly established cell-mass relation (Eq. 3), we demand that *z*(*t*) ∝ *m*(*t*) and *r*(λ) ∝ 1/(*C* + *D*). The first condition requires the concentration of proteins(s) synthesizing *L*, *z*/*m*, to be growth-rate independent. We analyzed the quantitative proteomics results across growth rates (**Supplementary Methods**) and found that the concentrations of hypothetical proteins involving in PCC formation (ZapA and FtsK)^9^ and the representative protein for septum formation (FtsZ)^34,35^ were relatively independent of the growth rate (coefficient of variation less than 10 %), while in comparison, a ribosomal protein RpsA increased linearly with growth rate (**b**). The specific rate *r* ∝ 1/(*C* + *D*) is a composite rate involving the serial events of chromosome replication, segregation, and cell division. We measured the *C*-period using a deep sequencing version of standard methods (**Supplementary Methods**). For a subset of 16 media that spanned growth rates from 0.4 h^−1^ to 1.7 h^−1^, the *C*-period remained roughly constant for fast growing cells while increasing significantly for slow growing cells. A simplifying relationship is revealed in (**c**), showing that *λC* (interpreted from the deep sequencing data, **Supplementary Methods**), and hence *λD*, is linearly related to *λ*(*C* + *D*) (interpreted from *ō*, **Supplementary Methods**). Thus, *r* could also be proportional to 1/*C* or 1/*D*.

**Extended Data Figure 8.**
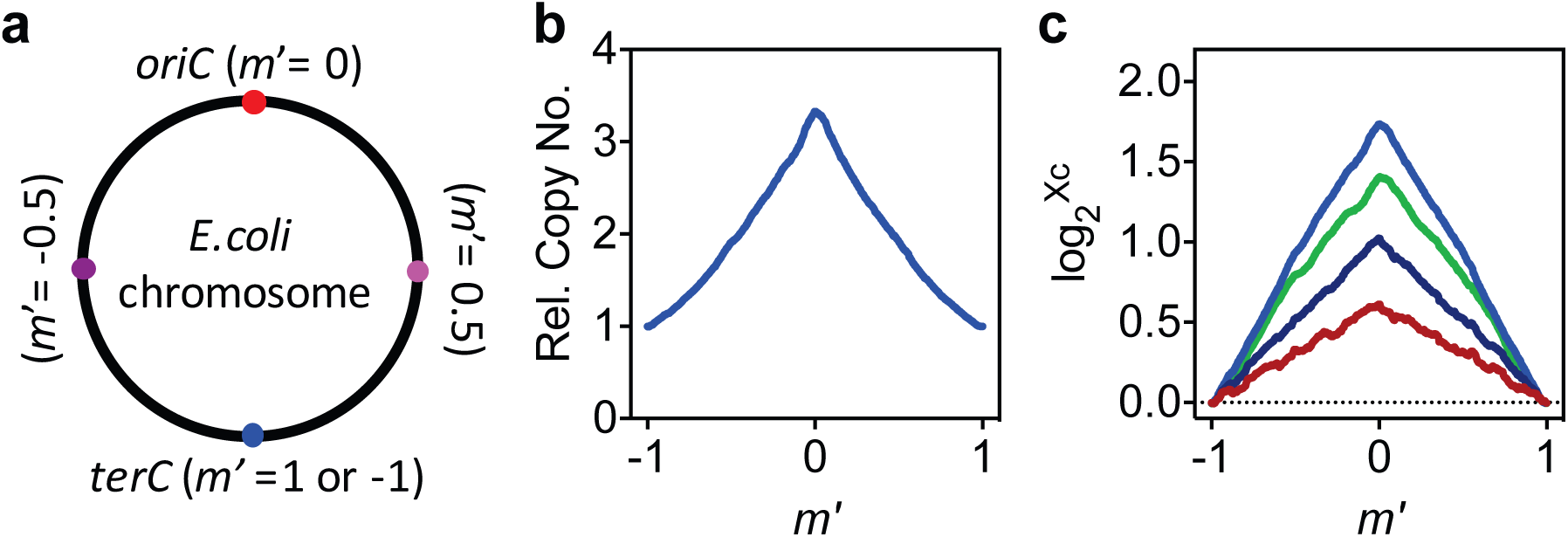
Measurement of the *C* period by deep sequencing. **a**, Definition of the relative chromosomal location (*m*′). To characterize the *C* period, the genome was binned into over 900 fragments of size 5,000bp. The relative chromosomal location for each fragment (*m*′) is defined by its relative location between *oriC* (*m*′ = 0) and *terC* (*m*′ = ±1). **b**, Dependence of relative gene copy number (*X*_*c*_) on chromosome location for cells grown in M1 (Extended Data Table 1). The relative gene copy number was obtained by normalizing the deep sequencing counts for each fragment to the count number for the fragment containing *terC* (**Supplementary Methods**). **c**, Linear correlation between the logarithm of the relative copy number of the fragment 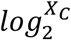 and *m*′. Different colors represent different growth media: blue, green, navy, and red correspond to M1, M3, M13, and M23, respectively (Extended Data Table 1).

**Extended Data Figure 9.**
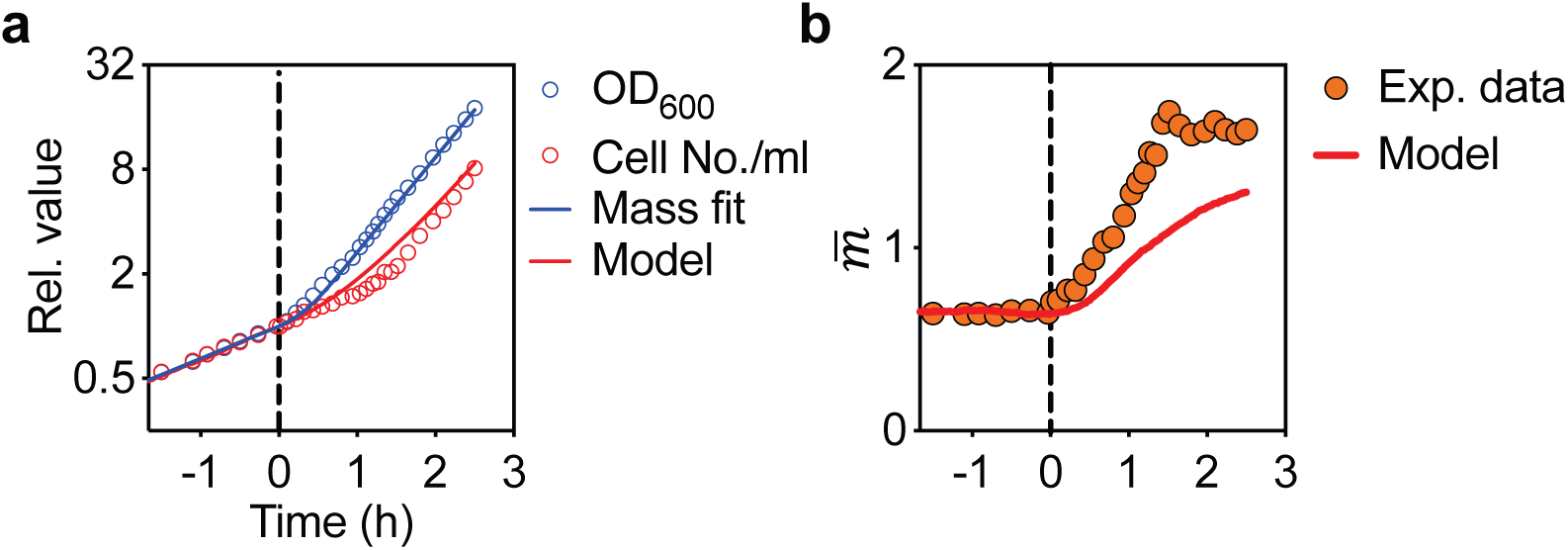
The integral threshold model is inadequate to produce the ‘rate maintenance’ phenomenon. **a**, Simulation of the model exhibits a delay in cell number growth rate during nutrition shift-up. A nutrition shift-up experiment was performed with a shift from growth medium M25 to M6 at time 0. Shown are the time course of OD_600_ (blue circles) and cell number (red circles). In the simulation, the total cell mass growth rate dynamics was described by a “ramp” fit to the transition in empirical mass growth rates. The cell number growth curve exhibited a pronounced shift relative to the mass growth curve, with a scale set by *C* + *D*, reminiscent of the rate-maintenance phenomenon. For details of the simulation, please see **Supplementary Model**. **b**, The model is insufficient to predict the dynamics of the 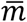 during the nutrition shift-up experiment. Experimental data of the 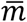 reach the new steady-state value faster than predicted by the model.

**Extended Data Table 1.**
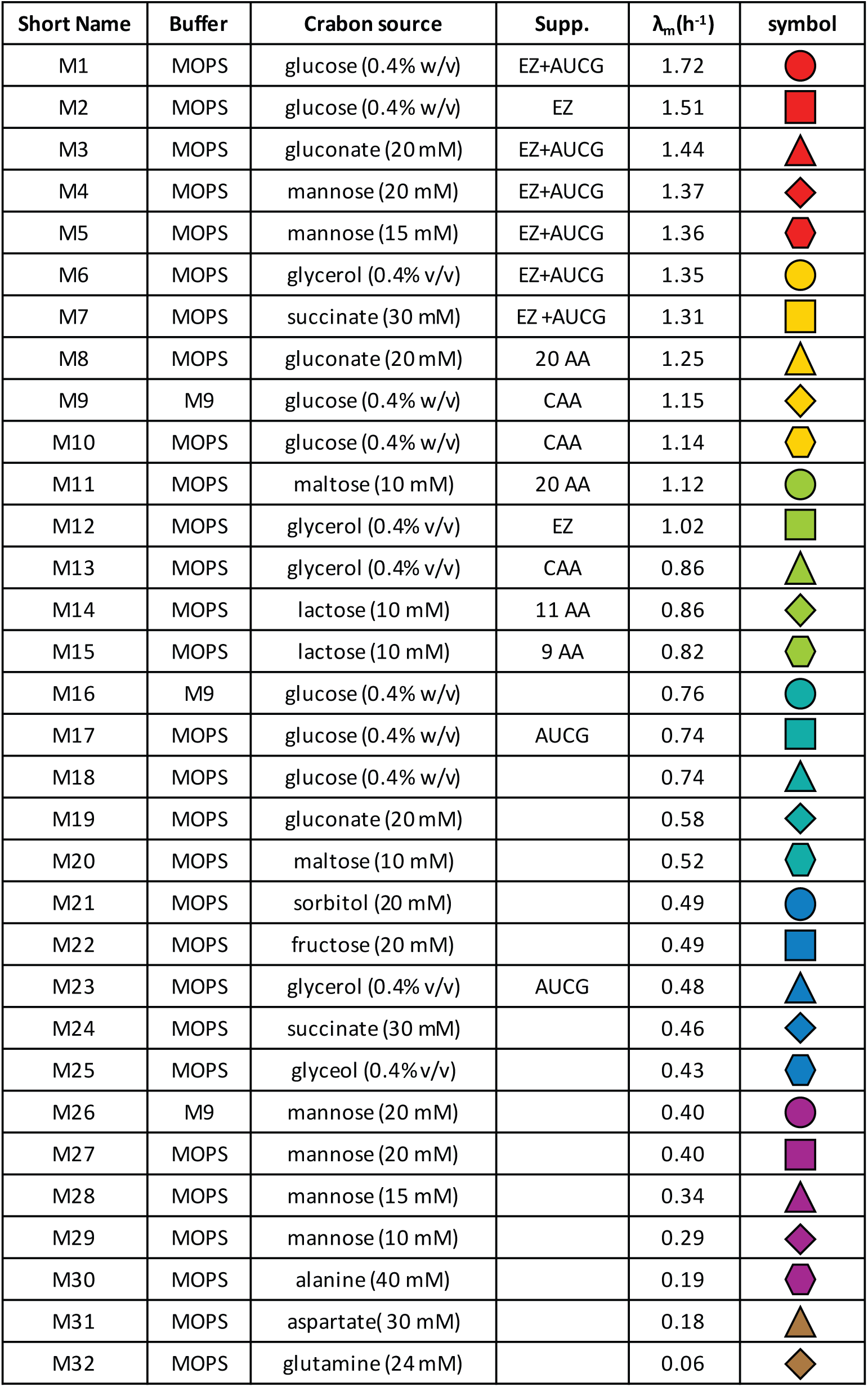

| Short Name | Buffer | Crabon source | Supp. | $\lambda_m(h^{-1})$ | symbol |
| --- | --- | --- | --- | --- | --- |
| M1 | MOPS | glucose (0.4% w/v) | EZ+AUCG | 1.72 |  |
| M2 | MOPS | glucose (0.4% w/v) | EZ | 1.51 |  |
| M3 | MOPS | gluconate (20 mM) | EZ+AUCG | 1.44 |  |
| M4 | MOPS | mannose (20 mM) | EZ+AUCG | 1.37 |  |
| M5 | MOPS | mannose (15 mM) | EZ+AUCG | 1.36 |  |
| M6 | MOPS | glycerol (0.4% v/v) | EZ+AUCG | 1.35 |  |
| M7 | MOPS | succinate (30 mM) | EZ +AUCG | 1.31 |  |
| M8 | MOPS | gluconate (20mM) | 20 AA | 1.25 |  |
| M9 | M9 | glucose (0.4% w/v) | CAA | 1.15 |  |
| M10 | MOPS | glucose (0.4% w/v) | CAA | 1.14 |  |
| M11 | MOPS | maltose (10 mM) | 20 AA | 1.12 |  |
| M12 | MOPS | glycerol (0.4% v/v) | EZ | 1.02 |  |
| M13 | MOPS | glycerol (0.4% v/v) | CAA | 0.86 |  |
| M14 | MOPS | lactose (10 mM) | 11 AA | 0.86 |  |
| M15 | MOPS | lactose (10 mM) | 9 AA | 0.82 |  |
| M16 | M9 | glucose (0.4% w/v) |  | 0.76 |  |
| M17 | MOPS | glucose (0.4% w/v) | AUCG | 0.74 |  |
| M18 | MOPS | glucose (0.4% w/v) |  | 0.74 |  |
| M19 | MOPS | gluconate (20 mM) |  | 0.58 |  |
| M20 | MOPS | maltose (10 mM) |  | 0.52 |  |
| M21 | MOPS | sorbitol (20 mM) |  | 0.49 |  |
| M22 | MOPS | fructose (20 mM) |  | 0.49 |  |
| M23 | MOPS | glycerol (0.4% v/v) | AUCG | 0.48 |  |
| M24 | MOPS | succinate (30 mM) |  | 0.46 |  |
| M25 | MOPS | glyceol (0.4%v/v) |  | 0.43 |  |
| M26 | M9 | mannose (20 mM) |  | 0.40 |  |
| M27 | MOPS | mannose (20 mM) |  | 0.40 |  |
| M28 | MOPS | mannose (15 mM) |  | 0.34 |  |
| M29 | MOPS | mannose (10 mM) |  | 0.29 |  |
| M30 | MOPS | alanine (40 mM) |  | 0.19 |  |
| M31 | MOPS | aspartate( 30 mM) |  | 0.18 |  |
| M32 | MOPS | glutamine (24 mM) |  | 0.06 |  |

**Extended Data Table 2.**

| Type of replicate | Biological Replicate |  |  |  |  |  | Cell number analyzed |
| --- | --- | --- | --- | --- | --- | --- | --- |
| Parameter<br>Medium | $\lambda, \bar{m}, \bar{o}, m_i$ | DW per cell | R+P per cell | Rel. FSC | Rel. Protein conc | C period | Cell volume |
| M1 | 9 | 2 | 2 | 5 | 3 | 3 | 761 |
| M2 | 4 | N.D. | N.D. | 3 | 3 | 3 | N.D. |
| M3 | 5 | 2 | 2 | 2 | 3 | 3 | N.D. |
| M4 | 6 | 2 | 2 | 3 | 3 | 3 | N.D. |
| M5 | 2 | 2 | 1 | 2 | N.D. | N.D. | N.D. |
| M6 | 7 | 2 | 2 | 4 | 3 | 3 | 1125 |
| M7 | 1 | N.D. | N.D. | 3 | N.D. | N.D. | N.D. |
| M8 | 2 | 2 | 2 | 2 | N.D. | N.D. | N.D. |
| M9 | 4 | 2 | 1 | 4 | N.D. | N.D. | 1448 |
| M10 | 9 | 2 | 2 | 5 | 3 | 3 | 832 |
| M11 | 1 | N.D. | N.D. | 1 | N.D. | N.D. | N.D. |
| M12 | 3 | N.D. | N.D. | N.D. | 3 | 3 | N.D. |
| M13 | 8 | N.D. | 2 | 5 | 3 | 3 | 2314 |
| M14 | 1 | N.D. | N.D. | 3 | N.D. | N.D. | N.D. |
| M15 | 1 | N.D. | N.D. | 3 | N.D. | N.D. | N.D. |
| M16 | 4 | 2 | 1 | 4 | N.D. | N.D. | 2465 |
| M17 | 3 | N.D. | N.D. | N.D. | 3 | 3 | N.D. |
| M18 | 9 | 2 | 2 | 5 | 3 | 1 | 1281 |
| M19 | 7 | 2 | 2 | 4 | 3 | 3 | 701 |
| M20 | 4 | 2 | 2 | 3 | N.D. | N.D. | 947 |
| M21 | 4 | 2 | 1 | 4 | N.D. | N.D. | 1121 |
| M22 | 6 | 1 | 1 | 3 | 3 | 1 | N.D. |
| M23 | 3 | N.D. | N.D. | N.D. | 3 | 1 | N.D. |
| M24 | 7 | 2 | 1 | 4 | 3 | 3 | 2129 |
| M25 | 9 | 2 | 1 | 5 | 3 | 3 | 1340 |
| M26 | 3 | 2 | 1 | 3 | N.D. | N.D. | 880 |
| M27 | 7 | 2 | 2 | 4 | 3 | 3 | 1014 |
| M28 | 6 | 2 | 2 | 3 | N.D. | N.D. | N.D. |
| M29 | 3 | 1 | 2 | 3 | N.D. | N.D. | 1868 |
| M30 | 3 | N.D. | N.D. | 3 | N.D. | N.D. | N.D. |
| M31 | 3 | N.D. | N.D. | N.D. | N.D. | N.D. | 2796 |
| M32 | 4 | N.D. | N.D. | N.D. | N.D. | N.D. | N.D. |

Note: The growth rate ( $\lambda$ ), population-averaged cellular mass $\text{OD}_{600} \cdot \text{ml}$ per $10^9$ cells ( $\bar{m}$ ), population-averaged cellular *oriC* No. ( $\bar{o}$ ), and the derived initiation mass ( $m_i$ ) were examined simultaneously for a same experimental culture, so these parameters have the same replicate number.

