## supporting information for "A general quantitative relation linking bacterial cell growth and the cell cycle"

### Supplementary methods

#### 1. Bacterial strain and growth media

The bacterial cell strain used in this study was *E. coli* K12 AMB1655, which was kindly provided by Antoine Danchin of AMAbiotics, Evry, France. Growth media with different nutrient composition were used to attain different growth rates. Detailed information on these media is listed in **Extended Data Table 1** and **Supplementary Table S1**.

#### 2. Culture procedure

Unless otherwise stated, we picked 3-5 single colonies from an LB agar plate and inoculated these colonies into 14 ml round bottom test tube containing 2 ml of the desired culture medium. Test tubes were kept in a shaker (220 RPM, Shanghai Zhichu Instrument Co., Ltd.) overnight (or longer, for slow growth media) at 37 °C as the seed culture procedure. Given the extremely slow growth rate for cells grown in media M29-M32, M18 was used as the growth medium for the seed culture procedure. The seed cultures were centrifuged and then washed with the desired culture media before inoculation into the desired medium for further experiments.

The following culture procedure was applied to demonstrate that exponential mass growth does not by itself indicate steady state growth. Seed cultures were inoculated into pre-warmed fresh medium with a starting $OD_{600}$ of approximately 0.02 and cultured in a water-bath shaker (150 RPM, Shanghai Zhichu Instrument Co., Ltd.) at 37 °C. The $OD_{600}$, cell number concentration, and population-averaged cellular origin number ($\bar{o}$), were characterized simultaneously at different time points to generate **Extended Data Fig. 1a-d**.

The following culture procedure was applied to verify the establishment of the steady state growth. Seed cultures were inoculated into pre-warmed fresh medium with a starting $OD_{600}$ of approximately 0.02 and cultured in a water-bath shaker (150 RPM) at 37 °C. Successive dilutions were carried out by transferring cell suspension into pre-warmed fresh medium with dilution ratio at 1:9 once the $OD_{600}$ reached 0.2 and repeated for another 2 rounds. Then the $OD_{600}$ and cell number concentration were characterized at different time points. During the sampling procedure, successive dilutions were carried out with dilution ratio 1:4 once the $OD_{600}$ reached 0.2. The time point for taking the initial sample is denoted as time 0 in **Extended Data Fig. 1f**.

Except for the above-mentioned protocols that were established for specific purposes, the following culture procedure was applied to establish steady state growth. Seed cultures were inoculated into pre-warmed fresh medium with a starting $OD_{600}$ of approximately 0.02 and cultured in a water-bath shaker (150 RPM) at 37 °C. Successive dilutions were carried out by transferring cell suspension into pre-warmed fresh medium with dilution ratio 1:9 once the $OD_{600}$ reached 0.2 and repeated for another 2 rounds. Then, the $OD_{600}$ growth curve was characterized to calculate the steady state growth rate ($\lambda$). Samples to quantify the $\overline{m}, C+D, C, \bar{o}$, dry weight per cell, RNA+Protein per cell, FSC per cell, etc., were collected at $OD_{600}$ approximately equal to 0.2.

#### 3. Dry weight per cell measurement

The dry weight per cell was inferred from the dry weight per $OD_{600}\cdot$ml, and cell number per $OD_{600}\cdot$ml. The dry weight per $OD_{600}\cdot$ml were characterized based on the method used by Erickson *et al*[1], and the cell number per $OD_{600}\cdot$ml were characterized by the flow cytometry as described below.

#### 4. Population-averaged $OD_{600}$·ml per $10^9$ cells characterization

When the $OD_{600}$ reached approximately 0.2, samples for characterizing $OD_{600}$·ml per $10^9$ cells were taken and the $OD_{600}$ carefully measured using a spectrometer (Genesys 10s, Thermo Fisher Scientific). To prepare sample for cell counting, an aliquot (see below) of cell suspension was immediately diluted with pre-cooled cell count buffer (0.9 % NaCl with 0.12% formaldehyde, filtered with 0.22 µm filter) and kept in an ice-water bath. The volume of cell suspension and cell count buffer was predetermined empirically based on the growth medium used. In order to get the most reliable cell counts, the cell concentration after this step should be within the range from $5\times10^6$ to $2\times10^7$ $ml^{-1}$.

Cell counting was performed with a flow cytometer equipped with 405 nm laser (CytoFLEX (S), Beckman Coulter Life Sciences). One aspect of this instrument that is essential for the current study was the ability to control the flow rate precisely. To 900 µl stain buffer (0.9 % NaCl with 1 µg $mL^{-1}$ DAPI, filtered with 0.22 µm filter), a 100 µl sample was added, then incubated in the ice-water bath for 3 minutes. The flow rate and running time were 1 µL $s^{-1}$ and 100 s, respectively. The trigger channel was set on SSC-H, the gain for FSC, SSC, and DAPI channel were set to 500, 500, and 2000 respectively. The DAPI-stained particles were deemed as the bacterial cells (**Fig. S1a-c)**. The $OD_{600}$·ml per $10^9$ cells was then calculated by dividing the $OD_{600}$ by the corresponding cell number concentration. Two different flow cytometry instruments, CytoFLEX or CytoFLEX S, were used in this study, and we confirmed that the calculated population-averaged $OD_{600}$·ml per $10^9$ cells was not related to the specific instrument applied.

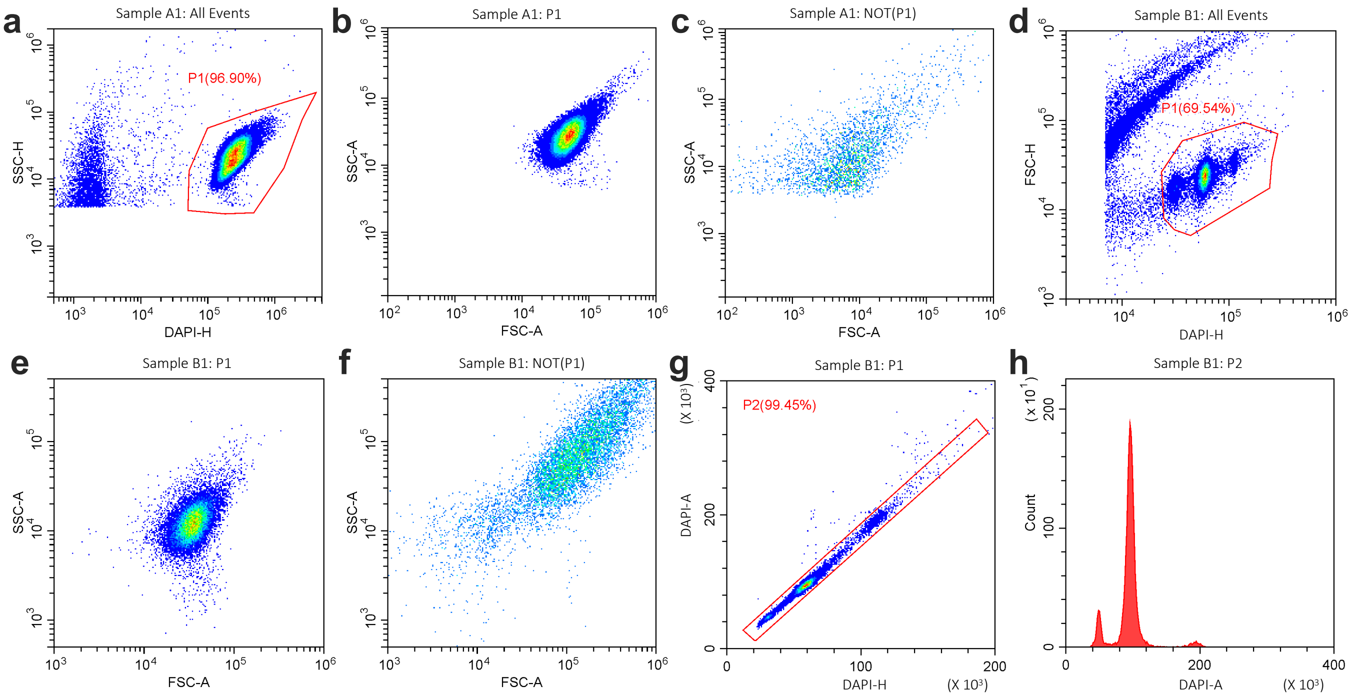

**Fig. S1**. Gating strategy used in flow cytometry analysis. **a-c**, Gating strategy used in cell counting and FSC characterization experiments. Particles in P1 in panel **a** were regarded as the bacterial cells. The FSC-A/SSC-A plots for P1 and Not P1 sub-populations are shown in panels **b** and **c**, respectively. **d-h**, Gating strategy used in experiments for population-averaged cellular oriC characterization. Particles in P1 in panel **d** were regarded as the bacterial cells. The FSC-A/SSC-A plots for P1 and Not P1 sub-populations are shown in panels **e** and **f**, respectively. A further gate was set on P1 to eliminate non-single-cell particles (**g**). **h**, The DAPI-A histogram for particles in P2 was used for calculating the population-averaged *oriC* number.

**5. Population-averaged cell volume characterization**

The population-averaged cell volume was characterized based on microscopic images as previously described[2]**.** Briefly, when the $OD_{600}$ of a steady-state culture reached approximately 0.2, 10 µl of the culture was immobilized by a 15 × 15 mm agarose pad (1% agarose prepared with 0.9% NaCl), and cover by a coverslip. Immobilized cells were imaged by a Nikon Ti-E microscope equipped with a 100× phase-contrast objective (N.A. = 1.45) and an Andor Zyla 4.2s. The images were acquired within 5 min at room temperature (RT) after immobilization. MicrobeTracker, a customized MATLAB (MathWorks)-based image-processing package was used to process the images. One pixel on the image equaled 0.065 µm, which was validated by a graticule. Approximately 1000 cells were analyzed to get the population-averaged cell volume. The exact number of cells analyzed for each growth condition have been listed in **Extended Data Table 1**.

**6. Characterization of the relative FSC of the cells**

The FSC-A of the cells were extracted from the experiments while characterizing the population-averaged $OD_{600}$·ml per $10^9$ cells. The FSC-A for cells grown in different growth media were normalized to the FSC-A for cells grown in M18. Only the FSC-A obtained by the same instrument have been used in **Fig. 1c** and Extended Data **Figure 2c**. We experienced difficulties in making effective comparisons of the FSC-A obtained by different instruments even at identical parameter settings.

**7. RNA and protein per cell measurement**

The RNA and protein per cell were inferred from the RNA per $OD_{600}$·ml , protein per $OD_{600}$·ml , and cell number per $OD_{600}$·ml . The RNA per $OD_{600}$·ml , and protein per $OD_{600}$·ml were characterized based on the method used by You *et al*[3], and the cell number per $OD_{600}$·ml were characterized by the flow cytometry as described above.

**8. Characterize the population-averaged cellular origin number ($\bar{o}$), $\lambda(C+D)$ and sum of *C* and *D* periods**

Replication run-out experiments were conducted for the quantification of $\bar{o}$ as previously described[2,4] with modification. Briefly, rifamycin and cephalexin were added into the cell suspension with final concentration at 300 µg/ml and 30 µg/ml, respectively, cells were allowed to grow for 2-3 mass doubling time to finish on-going replication. To 300 µl cell suspension after run-out experiments, 700 µl pre-cooled absolute ethanol was added by drops with continuous mixing. The fixed sample could be applied for further flow cytometer characterization after incubating for 30 minutes at 4 °C, or stored at 4 °C for at least 24 hours without affecting the results. To 20 µl fixed sample, 480 µl stain buffer (20 mM Tris·HCl, pH 8.0, 130 mM NaCl, 0.1% Triton X-100, 10 ng $mL^{-1}$ DAPI) were added directly, mixed well by vortex, and kept at room temperature for 3 minutes.

Flow cytometry analysis was performed using CytoFLEX (S) equipped with a 405-nm laser. The trigger channel was set on DAPI, and the gains for FSC, SSC, and DAPI channel were set to 500, 500, and 2000 respectively. $\bar{o}$ was calculated based on the distribution of the DAPI signaling of the run-out experiments (**Fig. S1d-h**). Two different flow cytometry instruments, CytoFLEX and CytoFLEX S, were used in this study, and we confirmed that the calculated $\bar{o}$ was not related to the specific instrument applied.

The $\lambda(C+D)$, and sum of $C$ and $D$ periods were evaluated from $\bar{o}$ using formula $\lambda(C+D) = ln(\bar{o})$, and $C+D = ln(\bar{o})/\lambda$ , respectively.

**9. Characterization of the $\lambda C$ and *C* period**

Theoretical analysis has revealed that, for a population of cells undergoing steady-state growth, the ratio of the frequencies of different genes depends on the *C* period, the location on the chromosome (m', **Extended Data Fig. 7**), and the mass doubling time (τ)[5]. The gene frequency ratio between *oriC* and *terC* has thus been widely used to quantify the *C* period. In a previous study, we developed a qPCR-based method to more precisely characterize the *C* period by taking more chromosomal sites into consideration. In this study, further improvements were made by taking the whole chromosome into consideration via deep sequencing.

Total DNA of steady state culture was extracted by using the genomic DNA purification kit (Tiangen) according to the manufacturer's protocol, with three DNA samples extracted from run-out cells as control experiments. DNA samples were sent to BGI for deep sequencing by an Illumina HiSeq 2000.

The genomic sequence was binned into over 900 fragments with 5000bp for each fragment. The m' coordinate was defined as the relative distance from the replication origin, with the distance from *oriC* to *terC* set at $\pm 1$. The total read count per fragment was found to be strongly biased by the GC content of the fragment for some samples, so the computeGCBias and correctGCBias programs were applied. We calculated the gene frequencies ($X_c$) for each fragment relative to that for *terC*. For run-out cells, $X_c$ was roughly equal to 1.0 for all m' (data not shown).

For each experimental sample, the correlation between $m'$ and $log_2^{X_c}$ was fit to two lines, depending on whether m' was greater or less than 0. The $\lambda C$, and *C* period were calculated as $\lambda C = |k| \times ln2$, and $C = |k| \times \tau$, respectively, where $k$ is the slope of the linear fit and $\tau$ is the mass doubling time. For each experimental sample, there are two different $k$ values of opposite sign. We noticed that they are approximately equal in absolute value. We took the mean value of the periods based on the individual $k$ as the *C* period for this experimental sample.

**10. Shift-up experiment**

For the shift-up experimental data presented in **Extended Data Fig. 8**, growth conditions were shifted from M25 to M6, corresponding to a growth rate shift from 0.43 to 1.35 $h^{-1}$. These two growth media have identical chemical composition except that M6 contains nucleotides (AUCG) and amino acids (EZ) but M25 does not. We thus avoided unnecessary manipulation, such as centrifugation or filtration, when conducting the shift, minimizing undesired perturbation on the cells other than the nutrition down-shift.

Before nutrition shift up, cells were growing in steady state in M25 as described in section 1.2. When $OD_{600}$ reached approximately 0.1, an aliquot of cell suspension was added to an equal volume of pre-warmed "2×M6" medium, which is M6 with extra AUCG and EZ. By doing so, the growth condition for the cells have been shifted from M25 to M6. The $OD_{600}$ and cell number concentration were followed before and after the shift up. Cells were diluted by the ratio 1:3 once the $OD_{600}$ reached 0.2. By taking the dilution ratio into account, continuously varying mass and cell number growth curves were produced.

**11. Proteomics**

To collect a sample for proteomics, a 15 ml cell suspension from steady-state culture at $OD_{600}$ approximately 0.2 was transferred into a 50 ml test tube, treated with liquid nitrogen immediately and then kept at -80 °C. After collecting all of the samples for 16 different growth media, these samples were thawed to 0 °C, centrifuged at 4 °C (10000g, 5 minutes), the supernatant discarded, and the pellet washed once with ice-water cooled PBS. After again discarding the supernatant, the pellets were stored at -80 °C.

The following procedure was carried out by Jingjie PTM Biolabs (Hangzhou, China). The sample was sonicated three times on ice using a high intensity ultrasonic processor (Scientz) in lysis buffer (8 M urea, 1% Triton X-100, 10 mM dithiothreitol, 1% Protease Inhibitor Cocktail, 2 mM EDTA). Total protein was extracted by trichloroacetic acid (TCA) methods and dissolved in urea solution (8 M). Protein concentration was quantified by BCA kit according to the manufacturer’s instructions.

For digestion, the protein solution was reduced with 5 mM dithiothreitol for 30 min at 56 °C and alkylated with 11 mM iodoacetamide for 15 min at room temperature in darkness. The protein sample was then diluted by adding 100 mM TEAB to urea concentration less than 2M. Finally, trypsin was added at 1:50 trypsin-to-protein mass ratio for the first digestion overnight and 1:100 trypsin-to-protein mass ratio for a second 4 h-digestion.

After trypsin digestion, peptide was desalted by Strata X C18 SPE column (Phenomenex) and vacuum-dried. Peptide was reconstituted in 0.5 M TEAB and processed according to the manufacturer’s protocol for Tandem Mass Tag (TMT) labeling kit. To conduct comparisons among 18 kinds of growth conditions (with 2 additional growth conditions which is not relate to current study) with three biological replicates for each condition, a mixed sample was prepared by mixing the all the 54 protein samples at equal protein amount. Protein samples were placed into 6 groups, with 9 different protein samples and the mixed sample in each group. Different TMT tags including 126, 127N, 127C, 128N, 128C, 129N, 129C, 130N, 130C, 131 were used for labeling different samples within the same group.

To improve proteome coverage, the tryptic peptides were fractionated by high pH reverse-phase HPLC using Agilent 300Extend C18 column (5 μm particles, 4.6 mm ID, 250 mm length). Briefly, peptides were first separated with a gradient of 8% to 32% acetonitrile (pH 9.0) over 60 min into 60 fractions. Then, the peptides were combined into 18 fractions and dried by vacuum centrifuging.

One fraction of the mixed peptides was dissolved in solvent A (0.1% formic acid in 2% acetonitrile), directly loaded onto a home-made reversed-phase analytical column (15-cm length, 75 μm i.d.). The gradient was comprised of an increase from 6% to 18% solvent B (0.1% formic acid in 90% acetonitrile) over 40 min, 18% to 28% in 12 min and climbing to 80% in 4 min then holding at 80% for the last 4 min, all at a constant flow rate of 300 nL/min on an EASY-nLC 1000 UPLC system.

The peptides were subjected to NSI source followed by tandem mass spectrometry (MS/MS) in Orbitrap Fusion Lumos (Thermo) coupled online to the UPLC. The electrospray voltage applied was 2.4 kV. The m/z scan range was 350 to 1550 for full scan, and intact peptides were detected in the Orbitrap at a resolution of 60,000. Peptides were then selected for MS/MS using NCE setting as 32 and the fragments were detected in the Orbitrap at a resolution of 30,000. Automatic gain control (AGC) was set at 5E4. Fixed first mass was set as 100 m/z.

The resulting MS/MS data were processed using Maxquant search engine (v.1.5.2.8). Tandem mass spectra were searched against SwissProt Escherichia coli K12 (MG1655) database (4313 sequences) concatenated with a reverse decoy database. Trypsin/P was specified as the cleavage enzyme allowing up to 2 missing cleavages. The mass tolerance for precursor ions was set at 20 ppm in First search and 5 ppm in Main search, and the mass tolerance for fragment ions was set at 0.02 Da. Carbamidomethyl on Cys was specified as fixed modification, and oxidation on Met was specified as variable modifications. FDR was adjusted to < 1% and minimum score for peptides was set > 40.

To convert the proteomics data into relative protein per $OD_{600}\cdot$ml, we characterized the total protein per $OD_{600}\cdot$ml for several different growth conditions. An empirical smooth line was used to describe the growth rate dependence of the total protein per $OD_{600}\cdot$ml and used for the conversion.

#### Supplementary Models

**Summary**

This text describes the models proposed in the main text, and the important features and biological relevance of them. We start by reviewing the classic understanding of coordination mechanism based on SMK growth law and the Donachie hypothesis (**Sect. 1)**. Next, we deduce the stochastic model from the linear relation (**Eq. 3**) described in the main text (**Sect. 2)**. After analyzing the features and drawbacks of the stochastic model (**Sect. 3**) we extend it to the integral-threshold model (**Sect. 4**). This model is then compared to the closest existing model (**Sect. 5**). In the last section (**Sect. 6**), the model is generalized to describe cell division during growth transitions, and the behaviors obtained are compared to the 'rate maintenance' phenomenon.

**1. Classic understanding of coordination**

Classic understanding of the coordination of cell mass growth, chromosome replication, and cell division in bacterial is manifested by the empirical growth law of Schaechter-Maaløe-Kjeldgaard (SMK)[6] and the constant initiation mass hypothesis by Donachie[7]. This framework states the followings for steady state cells in the fast-growth regime, i.e. doubling time shorter than 60 minutes:

1) average cell mass depends exponentially on growth rate[6];

2) the time interval *C*, from initiation to termination of one round of DNA replication, and *D*, from replication termination to corresponding cell division, sum to a constant[8];

3) the "initiation mass" (cellular mass per replication origin at the time of replication initiation) is constant[7].

Bremer *et al*[5], among others, have noted that the average number of origins per cell is

$$\bar{o} = 2^{\frac{C+D}{\tau}} = e^{\lambda(C+D)}. \quad \text{(S1)}$$

A growth rate dependent *C+D* results in a growth rate-dependent average mass per origin ($m_{po}$) as $m_{po} = \bar{m}(\lambda)/e^{\lambda(C+D)}$. The SMK growth law as reformulated by Donachie specifies that $\bar{m} \propto e^{\lambda(C+D)}$, and the growth rate dependence in $m_{po}$ cancels out. As the initiation mass $m_i$ is simply proportional to $m_{po}$, i.e. $m_i = m_{po}/\ln(2)$[9], the SMK growth law implies constant initiation mass, independent of growth rate. The above three statements support each other and form a consistent theoretical framework in the quantitative description of cell division.

**2. From the linear relation to the stochastic division model**

Our experiments have carefully examined the relation between average cell mass and growth rate, and revealed a new relation (**Eq. 3**). As the average cellular mass is the ratio between total cell mass and total cell number, the linear relation can be written as:

$$\frac{M(t)}{N(t)} = m_0 \lambda (C + D). \tag{S2}$$

$M(t)$ is the time-dependent total cell mass and $N(t)$ is the time-dependent total cell number. Then the above equation can be reformed to

$$\lambda N(t) = \frac{M(t)}{m_0(C + D)} = \frac{dN(t)}{dt}. \tag{S3}$$

We interpreted this population level equation as resulting from a single cell division process with division rate $k(t)$:

$$k(t) = \frac{m(t)}{m_0(C + D)}. \tag{S4}$$

where $dN(t)/dt$ is replaced by cell division rate $k$ and $M(t)$ is replaced by cellular mass $m(t)$. The sum of all cells in a system with different cell mass $m_j(t)$ gives the population level cell number growth rate $\frac{dN(t)}{dt} = \sum \frac{m_j(t)}{m_0(C+D)} = \frac{\sum m_j(t)}{m_0(C+D)} = \frac{M(t)}{m_0(C+D)}$, so that the population level relation is preserved.

**3. The stochastic division model**

Inspired by the above single cell division rate, we propose a stochastic cell division model based on the following assumptions:

(1) The division rate of individual cell follows **Eq. S4**

(2) Cellular mass $m(t)$ increases exponentially from birth mass $m_b$ with rate $\lambda$,

$$m(t) = m_b e^{\lambda t} \tag{S5}$$

(3) *C+D* is growth rate-dependent in accordance with experimental observations (**Eq. 2** and **Extended Data Fig. 3**) with fitted parameters $\alpha = 0.27, \beta = 0.99$,

(4) Cell mass is evenly distributed between daughter cells upon division.

This model considers cell division as a stochastic process, where the "hazard rate" $k(t)$ of a division event depends on the cellular mass at the current time. I.e. the division process is a non-homogeneous Poisson process[10].

The hazard rate is by definition[10] the conditional probability that the time of the cell division event $X$ (alternatively, cell age at division) lies in the interval $(t, t + dt)$, given that the cell has survived without dividing to time $t$: $k(t) \equiv P\{X \in (t, t + dt) | X > t, m_b\}$. We can get the probability density of cell division $p(t)$ defined as the probability of a cell of birth mass $m_b$ dividing at time $t$,

$$p(t) \equiv P\{X \in (t, t + dt) \mid m_b\} = k(t) e^{-\int_0^t k(s) ds} \tag{S6}$$

Integrating the cell division probability function $p(t)$ over time yields the cumulative distribution of cell division $F(t)$ that measures the probability of a cell to divide at a time between birth and age $t$,

$$F(t) = \int_0^t p(t')dt' = \int_0^t k(t')e^{-\int_0^{t'} k(s)ds}dt' \quad \text{(S7)}$$

Both quantities are plotted in Fig. S2.

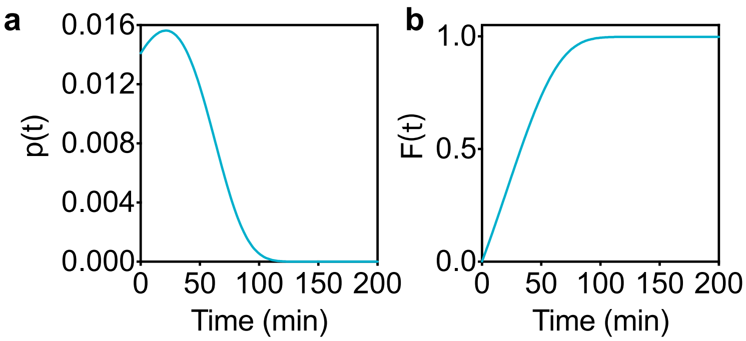

**Fig. S2**. Probability density function $p(t)$ (**a**) and cumulative distribution function $F(t)$ (**b**) of the stochastic division model.

Knowing the probability of cell division, we are able to obtain the expected cell mass at division given that a cell is born with mass $m_b$:

$$\langle m_f | m_b \rangle = \int_0^\infty m(t)\ P\{X \in (t, t+dt) \mid m_b\}\, dt. \quad \text{(S8)}$$

We can use the substitution $u = e^{\lambda t} - 1$, to derive the expected cell division mass

$$\langle m_f | m_b \rangle = m_0 \lambda (C + D) + m_b. \quad \text{(S9)}$$

The expected mass added between birth and division, $\Delta m \equiv \langle m_f | m_b \rangle - m_b = m_0 \lambda (C + D)$, is thus independent of the birth mass. This shows that the cellular mass-dependent stochastic division model that we have proposed in this paper is an exact adder model.

Though successful in its various predictions (Fig. 4b-d), the stochastic division model is inadequate to describe the distribution of division mass, nor inter-division time (**Extended Data Fig. 5**). Compared to the experimental data of Wallden *et al.*[11], the predicted distributions fail to capture the relatively narrow peaks in the empirical distributions (**Fig. S3**). The large portion of small cells and cells with short interdivision time is the main problem.

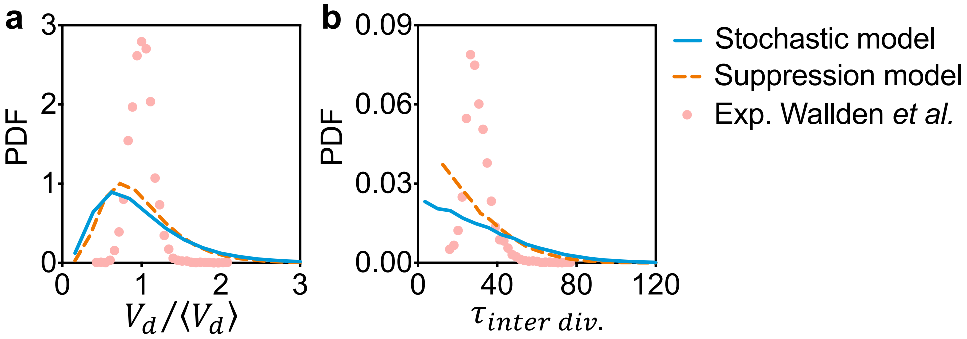

**Fig. S3**. Normalized division mass distribution (**a**) and inter-division time distribution (**b**) of the stochastic model (blue lines) and the suppression model (orange dashed lines) compared to the experimental data of Wallden *et al.*[11] (red circles). Simulations were performed in a mother machine mode (only one cell was kept after division) where $10^5$ cells were simulated with uniform initial cell mass.

The events with short inter-division times is due to the non-zero division probability at small time $t$, i.e., shortly after a division; see **Fig. S2 (a)**. At cell birth, the division probability density function (**Eq. S6**) $p(t=0) = k(0)e^{-\int_0^0 k(s)ds} \neq 0$ is determined by $k(0) = \frac{m_b}{m_0(C+D)}$, as the exponential $e^{-\int_0^0 k(s)ds}$ equals 1. The relatively large value of division probability $p(t=0) = \frac{m_b}{m_0(C+D)}$ gives rise to the large portion of cells that divide immediately after birth.

In attempt to fix this problem, we first introduced a refractory period after each division event. In this way, the division rate and division probability density function are suppressed around $t = 0$ (**Fig. S4**). However, the simulation of this suppression model shows almost the same wide distribution of division mass and inter-division time (see **Fig. S3** orange dashed lines). I.e., this modification is not a solution.

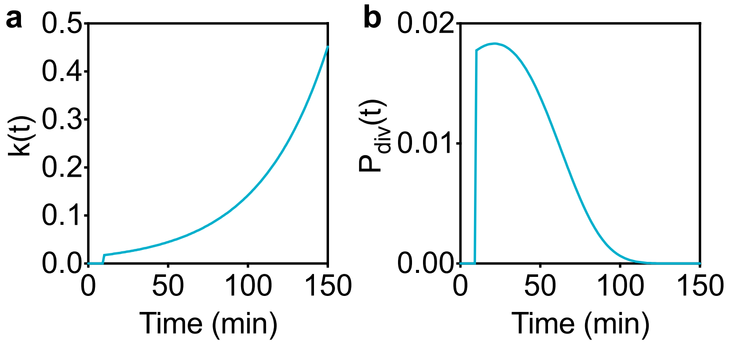

**Fig. S4**. Division rate and probability density function of the modified model with a refractory period of 10 mins for cells of doubling time of 30 mins as an example.

As the division rate $k(t)$ is nonzero at $t = 0$ while its integral $\int_0^t k(s)ds$ is zero at $t = 0$, let us turn our attention to the latter, denoted as

$$L(t) \equiv \int_0^t k(s)ds\,. \tag{S10}$$

Biologically, this integral, corresponding to **Eq. 5** of the main text, may describe the accumulation of a certain "licensing product" as discussed in the main text and **Extended Data Fig. 6**.

The cumulative distribution function $F$ of the stochastic model can be written as an explicit function of $L(t)$,

$$F\big(L(t)\big) = \int_0^t p(t')dt' = \int_0^t \frac{dL(t')}{dt'} e^{-L(t')}dt' = 1 - e^{-L(t)}. \tag{S11}$$

This relationship is plotted as the blue curve in the following figure.

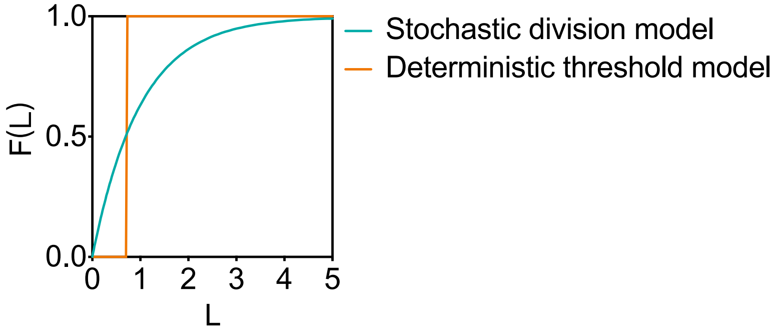

**Fig. S5**. Cumulative distribution $F(L)$ as a function of cell mass integral $L$. Both the stochastic model and deterministic model are plotted.

##### 4. The integral threshold model

To better address the cell-size distribution problem of the stochastic model, we turn to the cumulative distribution $F(L)$. The problem of consecutive cell division due to a finite probability $p(t \sim 0)$ is manifested as a finite slope for $F(L)$ near $L = 0$. To suppress consecutive cell division, we can consider an alternative form of $F(L)$, e.g., a step function in the simplest case (see **Fig. S5** orange line). In this case, the cell division is no longer stochastic but deterministic.

A step function of $F(L)$ has the following mathematical form,

$$F\big(L(t)\big) = \begin{cases} 1 & L(t) \geq L_c \\ 0 & L(t) < L_c \end{cases} \tag{S12}$$

which indicates that a cell divides when the accumulation of some division licensing product $L(t)$ reaches a threshold $L_c$. We refer to this as the integral-threshold model.

It is known that the population-averaged cellular mass $\overline{m}$ is proportional to population-averaged division mass $m_d$ with a factor of ln2 in a steady state batch culture $\overline{m} = \ln 2 \cdot m_d$[12]. Fitting it to the expected average cell mass $m_0 \lambda (C + D)$, we get the value $L_c = 0.722$. We observe that cells starting from different random initiation masses converge to the same cellular mass dynamics around the expected average cell mass (see **Fig. S6**). The linear relation (**Eq. 3**) is also reproduced with the same constant $L_c$ for all growth rates (**Fig. S7**).

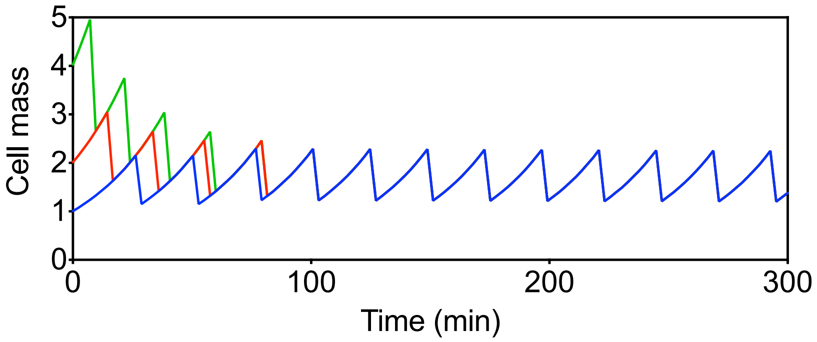

**Fig. S6**. Simulation of single cells having different initial cell masses ($\mathrm{m_b} = 1, 2, 4$) in the integral threshold model. Doubling time is 24 mins.

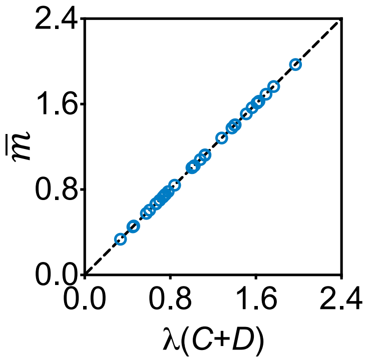

**Fig. S7**. Average cell mass calculated by $\bar{m} = \ln 2 \cdot m_d$ plotted with respect to $\lambda(C + D)$, where $m_d$ is the division mass of steady state single cells in the simulations. The dashed line represents the linear relation **Eq. 3**.

A purely deterministic model cannot produce a distribution of division masses. To compare to the experimentally observed distribution of division mass and inter-division time, we introduce stochastic fluctuations during the accumulation of $L$,

$$L(\mathrm{t}) = \int_0^t \big(k(s) + \eta(s)\big)ds. \tag{S13}$$

Here $\eta(s) \propto m(s)$ is taken to be a cellular mass dependent Gaussian noise.

With a relative noise level of ~$10\ \%$ and threshold $L_c = 0.685$, we are able to get distributions resembling the observed ones for both division mass and inter-division time (see **Extended data Fig. 6**).

**5. Comparison to existing models of cell division**

Most existing models about cell division were built on the classic framework presented in Sec.1. For example, the models proposed by Ho *et al*.[13], and Micali *et al*.[14,15] use the assumption of constant initiation mass. One can save these models by applying the experimentally measured growth rate dependent initiation mass (**Eq. 4**). But the complex form of the growth rate dependency complicates the biological interpretation.

There is also a class of threshold model not relying on initiation mass assumptions as proposed theoretically by Ghusinga *et al*.[16] and used by Si *et al.*[17] in their study of oscillatory expressed FtsZ cells. Si *et al*'s model, based on their study of oscillatory expressed cell cycle control genes[17], is a threshold model on the abundance of cell division proteins FtsZ. Cell division occurs when the amount of FtsZ per cell, denoted as $X(t)$, passes a threshold $X_0$. In the above framework, this amounts to the following form for the cumulative distribution function:

$$F\big(X(t)\big) = \begin{cases} 1 & X(t) \geq X_0 \\ 0 & X(t) < X_0 \end{cases} \tag{S14}$$

The biological meaning of the model with **Eq. S14** is quite different from that of our model (**Eq. S12**), as our model emphasizes the licensing product $L(t)$ which is synthesized by proteins such as FtsZ, whereas the model of Si *et al.* directly applies the threshold on these proteins.

For cells under balanced growth, the model of Si *et al*. can be rearranged to a closer form to ours to facilitate finer comparisons. Assuming that cellular proteins including FtsZ have constant concentrations throughout the cell cycle, we have in this case $[\mathrm{FtsZ}] \equiv X(t)/m(t)$. We can thus write

$$X(t) = \int_0^t \frac{\lambda m(s)}{[\mathrm{FtsZ}]} ds + X(t=0) \tag{S15}$$

since $m(t) = \lambda \int_0^t m(s)ds + m(t=0)$ for exponentially growing cells. Comparing **Eq. S15** to **Eq. 5** of our model, we see that the two models are mathematically the same, except for 2 differences: 1) the accumulation rate is the cell mass growth rate λ while it is $1/(C+D)$ in our model; 2) $X(t)$ is reset to its birth value $X(t=0)$ instead of 0 in our model. The first difference makes the two models biologically distinct: In the model of Si *et al.*, FtsZ protein and its product (the Z-ring) are interchangeable, whereas in our model the two are manifestly distinct. Regarding the second difference, the definition of $L$ can be easily modified to include a nonzero reset value, while yielding the same division behavior.

Both the models of Si *et al.* and ours consider cell division as a mass-dependent process instead of licensed by DNA replication events. Si *et al.*[17] do not attempt to model the growth rate-dependence of the average cell mass, which remains constant if the concentration $[\mathrm{FtsZ}]$ and the threshold $X_0$ are growth rate-independent. These authors also completely deny the effect of DNA replication initiation on cell division. This effect is kept in our model as the specific rate of license synthesis, as $1/(C+D)$, $1/C$, or $1/D$. As this rate is growth rate-dependent, our model is able to capture the cell division process for all growth rates without changing any parameters. The simulation results show that the linear relation between average cell mass and growth rate is well-captured (**Fig. 4b**).

**6. Cell division during growth transition**

As both the stochastic model and the integral threshold model can correctly predict the average cellular mass in steady state, we next extend these models to check how they behave during growth transitions.

'Rate maintenance' is an important phenomenon that occurs when bacteria are transferred from a slow-growth medium to a fast-growth medium[18]. It is observed that while the rate of cell mass growth changes rapidly to the rate of post-shift growth, the rate of cell number growth is *maintained* at that of pre-shift growth for certain period. This phenomenon is often considered as strongly supporting a link between chromosome replication and cell division[19].

For both the stochastic division model as defined in **Sec.3** and the integral threshold model as defined in **Sec.4**, we take the mass growth rate $\lambda_m(t)$ to be a linear fit of the observed transition during nutrition shift-up from growth media M25 to M6 (**Fig. S8a**). The time-dependent C+D is then derivable from **Eq. 2** (red curve in **Fig. S8a**). Both models are first simulated at the pre-shift growth condition until a steady state mass distribution was achieved. Then by random sampling from this mass distribution as the initial condition, we restart the simulation with pre-shift growth rate and perform growth rate transition as described above.

For both the stochastic division model and the integral-threshold model, the cell number growth curve exhibits a pronounced horizontal shift relative to the mass growth curve, reminiscent of the rate-maintenance phenomenon (**Fig. S8**). However, it is difficult to fully capture the instantaneous cell number growth rate (**Fig. S8c**) and the average cell mass data (**Fig. S8d**). Instead of a firm 'maintenance' of cell number growth rate, our models predict a smooth transition of cell number growth rate after medium shift.

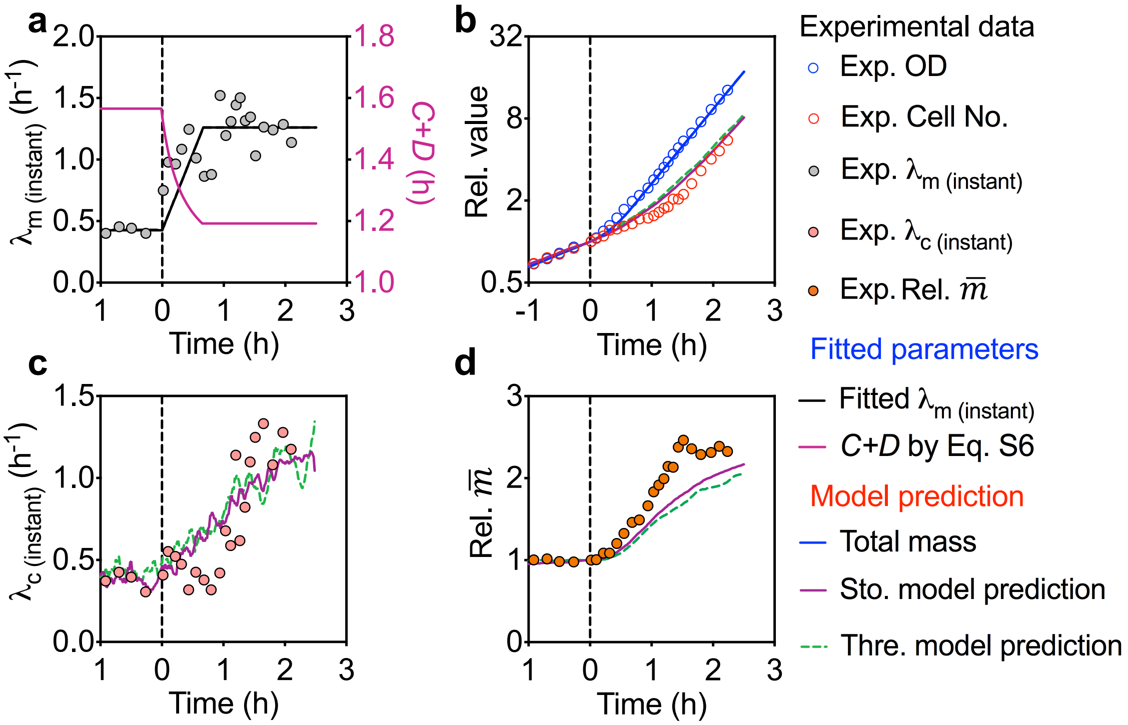

**Fig. S8**. Simulation of both the stochastic model and integral threshold model show delay in cell number growth rate during nutrition shift-up experiment. **a**. Nutrition shift-up experiment was performed with a shift in growth media from M25 to M6 at time 0 while following $OD_{600}$ (blue circles) and cell number (red circles). In the simulation, the total cell mass growth rate dynamics was fitted according to a "ramp" fit to the transition in empirical mass growth rates. The dynamics of the growth rate during transition was fit by a simple linear function connecting the two time-invariant steady state growth rates, using a least-squared fit of the time while the time-dependent value of *C+D* was calculated by **Eq. 2** (purple line). **b**. The cell number growth curve exhibits a pronounced shift relative to the mass growth curve, with a scale set by *C+D*, reminiscent of the rate-maintenance phenomenon. **c**. The model is insufficient to predict the dynamics of cell number growth rate as both models predict smoothly increasing growth rate $\lambda_c$, while the experimental growth rate shifts during a short period from one state to another. **d**. The experimental data shows that $\overline{m}$ reaches the new steady-state value faster than predicted by the model.

### Supplementary Table

#### 1. Chemical composition of the buffer and supplements for growth media

| | Chemical compound | Final conc. | Notes |
|---|---|---|---|
| MOPS Buffer (PH 7.4) | MOPS | 40 mM | Adjust to PH=7.4 |
| | Tricine | 4 mM | |
| | $FeSO_4$ | 10 µM | |
| | $NH_4Cl$ | 9.5 mM | |
| | $K_2SO_4$ | 0.276 mM | |
| | $CaCl_2$ | 0.5 µM | |
| | $MgCl_2$ | 0.525 mM | |
| | NaCl | 50 mM | |
| | $(NH_4)_6Mo_7O_{24}$ | 3 nM | Micronutrient |
| | $H_3BO_3$ | 0.4 µM | Micronutrient |
| | $CoCl_2$ | 30 nM | Micronutrient |
| | $CuSO_4$ | 10 nM | Micronutrient |
| | $MnCl_2$ | 80 nM | Micronutrient |
| | $ZnSO_4$ | 10 nM | Micronutrient |
| | $K_2HPO_4$ | 1.32 mM | |
| M9 Buffer | $Na_2HPO_4$ | 47 mM | |
| | $KH_2PO_4$ | 22 mM | |
| | NaCl | 8.5 mM | |
| | $NH_4Cl$ | 18.7 mM | |
| | Thiamine HCl | 1 mM | |
| | $MgSO_4$ | 2 mM | |
| | $CaCl_2$ | 0.1 mM | |
| EZ | Alanine | 0.8 mM | |
| | Arginine | 5.2 mM | |
| | Asparagine | 0.4 mM | |
| | Aspartic acid | 0.4 mM | |
| | Cysteine | 0.1 mM | |
| | Glutamic acid | 0.6 mM | |
| | Glutamine | 0.6 mM | |
| | Glycine | 0.8 mM | |
| | Histidine | 0.2 mM | |
| | Isoleucine | 0.4 mM | |
| | Leucine | 0.8 mM | |
| | Lysine | 0.4 mM | |
| | Methionine | 0.2 mM | |
| | Phenylalanine | 0.4 mM | |
| | Proline | 0.4 mM | |
| | Serine | 10 mM | |
| | Threonine | 0.4 mM | |
| | Tryptophan | 0.1 mM | |
| | Tyrosine | 0.2 mM | |
| | Valine | 0.6 mM | |
| | Thiamine HCl | 10 µM | Vitamin |
| | Calcium pantothenate | 10 µM | Vitamin |

| | | | |
|---|---|---|---|
| | *p*-aminobenzoic acid | 10 µM | Vitamin |
| | *p*-hydroxybenzoic acid | 10 µM | Vitamin |
| | 2,3-dihydroxybenzoic acid | 10 µM | Vitamin |
| 20 AA | Alanine | 0.8 mM | |
| | Arginine | 5.2 mM | |
| | Asparagine | 0.4 mM | |
| | Aspartic acid | 0.4 mM | |
| | Cysteine | 0.1 mM | |
| | Glutamic acid | 0.6 mM | |
| | Glutamine | 0.6 mM | |
| | Glycine | 0.8 mM | |
| | Histidine | 0.2 mM | |
| | Isoleucine | 0.4 mM | |
| | Leucine | 0.8 mM | |
| | Lysine | 0.4 mM | |
| | Methionine | 0.2 mM | |
| | Phenylalanine | 0.4 mM | |
| | Proline | 0.4 mM | |
| | Serine | 10 mM | |
| | Threonine | 0.4 mM | |
| | Tryptophan | 0.1 mM | |
| | Tyrosine | 0.2 mM | |
| | Valine | 0.6 mM | |
| 11 AA | Alanine | 0.8 mM | |
| | Arginine | 5.2 mM | |
| | Asparagine | 0.4 mM | |
| | Aspartic acid | 0.4 mM | |
| | Glutamic acid | 0.6 mM | |
| | Glutamine | 0.6 mM | |
| | Glycine | 0.8 mM | |
| | Proline | 0.4 mM | |
| | Serine | 10 mM | |
| | Threonine | 0.4 mM | |
| | Cysteine | 0.1 mM | |
| 9AA | Histidine | 0.2 mM | |
| | Isoleucine | 0.4 mM | |
| | Leucine | 0.8 mM | |
| | Tryptophan | 0.1 mM | |
| | Valine | 0.6 mM | |
| | Lysine | 0.4 mM | |
| | Methionine | 0.2 mM | |
| | Phenylalanine | 0.4 mM | |
| | Tyrosine | 0.2 mM | |
| CAA | Casamino acids | 0.2% (w/v) | |
| AUCG | Adenine | 0.2 mM | |
| | Cytosine | 0.2 mM | |
| | Uracil | 0.2 mM | |
| | Guanine | 0.2 mM | |

**Remarks:**

The composition of the MOPS buffer, EZ, and AUCG are based on Neidhardt Supplemented MOPS Defined Medium, prepared as described on the University of Wisconsin-Madison *E.coli* Genome Project website. The concentration for each amino acid used in 20 AA, 11 AA, 9 AA is identical to that used in EZ. The composition of M9 buffer and CAA are kept in accordance with the protocol presented on the OpenWetWare website.
